# Machine Learning Driven Optimization for High Precision Cellular Droplet Bioprinting

**DOI:** 10.1101/2024.09.04.611131

**Authors:** Jaemyung Shin, Minseok Kang, Kinam Hyun, Zhangkang Li, Hitendra Kumar, Kangsoo Kim, Simon S. Park, Keekyoung Kim

## Abstract

Controlled volume microliter cell-laden droplet bioprinting is important for precise biologics deposition, reliably replicating 3D microtissue environments for building cell aggregates or organoids. To achieve this, we propose an innovative machine-learning approach to predict cell-laden droplet volumes according to input parameters. We developed a novel bioprinting platform capable of collecting high-throughput droplet images and generating an extensive dataset for training machine learning and deep learning algorithms. Our research compared the performance of three machine learning and two deep learning algorithms that predict droplet volume based on numerous bioprinting parameters. By adjusting bioink viscosity, nozzle size, printing time, printing pressure, and cell concentration as input parameters, we precisely could control droplet sizes, ranging from 0.1 µL to 50 µL in volume. We utilized a hydrogel precursor composed of 5% gelatin methacrylate and a mixture of 0.5% and 1% alginate, respectively. Additionally, we optimized the cell bioprinting process using green fluorescent protein-tagged 3T3 fibroblast cells. These models demonstrated superior predictive accuracy and revealed the interrelationships among parameters while taking minimal time for training and testing. This method promises to advance the mass production of organoids and microtissues with precise volume control for various biomedical applications.

## Introduction

The bioprinting has the potential to replace two-dimensional cell cultures by more closely replicating the three-dimensional (3D) microtissue environment ^1,2^. Moreover, its versatility enables a broad spectrum of applications, spanning tissue engineering and regenerative medicine ^3,4^, transplantation ^5^, pharmaceuticals and high-throughput screening ^6^, and cancer research ^7^. Among many types of bioprinting, droplet bioprinting offers significant advantages in cell-laden droplet bioprinting, leveraging its inherent simplicity and agility. In recent years, there has been a significant increase in research both scaffold-free of stem cell-derived organoids and cell-laden bioprinting ^8–10^. For scaffold-free bioprinting, Lawlor *et al.* bioprinted kidney organoids and conducted a comparative analysis with manually created organoids to assess the potential of bioprinting technology ^11^, while similarly, Shin *et al.* developed a cost-effective, customized 3D bioprinter to produce 2 µL droplets containing 8 × 10³ nephron progenitor cells to make kidney organoids ^12^. For cell-laden bioprinting, Sakthivel *et al.* utilized inkjet bioprinting to fabricate a cell-laden gelatin methacrylate (GelMA) microgel array on an elastic composite substrate that was periodically stretched, demonstrating the positive effects of dynamic stretching ^13^.

Despite its promising potential, droplet bioprinting faces significant challenges, particularly in the intricate process of controlling and optimizing numerous bioprinting parameters such as temperature, bioink properties, printing time, printing speed, nozzle size, and dispensing pressure, etc. ^14^. Overcoming these hurdles traditionally requires extensive experimentation with substantial human resources and quantities of bioink materials. This challenge is further amplified when working with stem cells, which are notably more sensitive than other cell sources such as immortalized cell lines and primary cells ^15^. The use of stem cells underscores the critical importance of achieving precise control over droplet volume, as even minor variations can significantly impact cell viability, functionality, and degree of maturity ^16^. The quality of bioprinted outcomes is significantly influenced by the printing parameters ^17^. Minor changes within various printing parameters can alter the printed outcome ^18^. Hence, Webb *et al.* have developed a generalized evaluation method, termed Parameter Optimization Index, for assessing bioprinted samples. This method utilizes various bioprinting parameters, including biomaterial composition, nozzle size, printing speed, and printing pressure ^19^. In addition, James *et al.* investigated the relationships between changes in mechanical properties, degradation, and swelling ratio by optimizing the hydrogel composition parameters ^20^. They found that the printing parameters were primarily influenced by the composition of the hydrogels used.

The combination of machine learning (ML) with advanced bioprinting technology can potentially accelerate research in tissue engineering and regenerative medicine ^21^. ML refers to computer programs that, based on big data, autonomously learn to predict the future or make decisions ^22^. It is an artificial intelligence paradigm that goes beyond simple initial data training, continuously collecting and learning from data to enhance accuracy. In the development principles of ML, there are three main paradigms: supervised learning, which involves training on labeled datasets; unsupervised learning, which processes unlabeled data to discover hidden patterns or structures within the dataset; and reinforcement learning, where algorithms learn through feedback in the form of rewards or penalties (**Fig. 1a**) ^23^. DL is a subset of ML that is based on artificial neural networks, which are similar to the human nervous system (**Fig.1b**) ^24^. One key difference between ML and DL is that DL can collect and process data without undergoing data preprocessing tasks typically required in ML (**Supplementary Table S1**) ^25,26^. Verheyen *et al.* used supervised ML to construct predictive frameworks with material databases and assessed the predictability of soft granular material design spaces ^27^. ML can identify patterns within given data and develop models through training ^28^. The algorithm predicts the optimal parameter combination in the developed user interface webpage by considering the relationship between input parameters and the corresponding droplet volume outcomes. To provide users with enhanced controllability and printability, we established a high-throughput bioprinting platform incorporating data-driven ML and deep learning (DL) for bioprinting the array of microtissues or organoids. We developed an customized 3D bioprinter with a high throughput droplet image acquisition system for the thousands of printed droplet based on various bioprinting parameters (e.g., bioink viscosity, nozzle size, printing time, printing pressure, and cell concentration). We also developed software to automatically measure droplet volume using image processing and transfered the data into the ML algorithms. The combination of ML with advanced bioprinting technology can potentially accelerate research in tissue engineering and regenerative medicine ^21^. In addition, the approach utilizing ML and DL to optimize bioprinting parameters will significantly reduce human time and effort in bioprinting preparation ^29^. Finally, our ML and DL-integrated bioprinting system promises to streamline intricate printing procedures, leading to swifter and more effective outcomes.

**Fig. 1:**
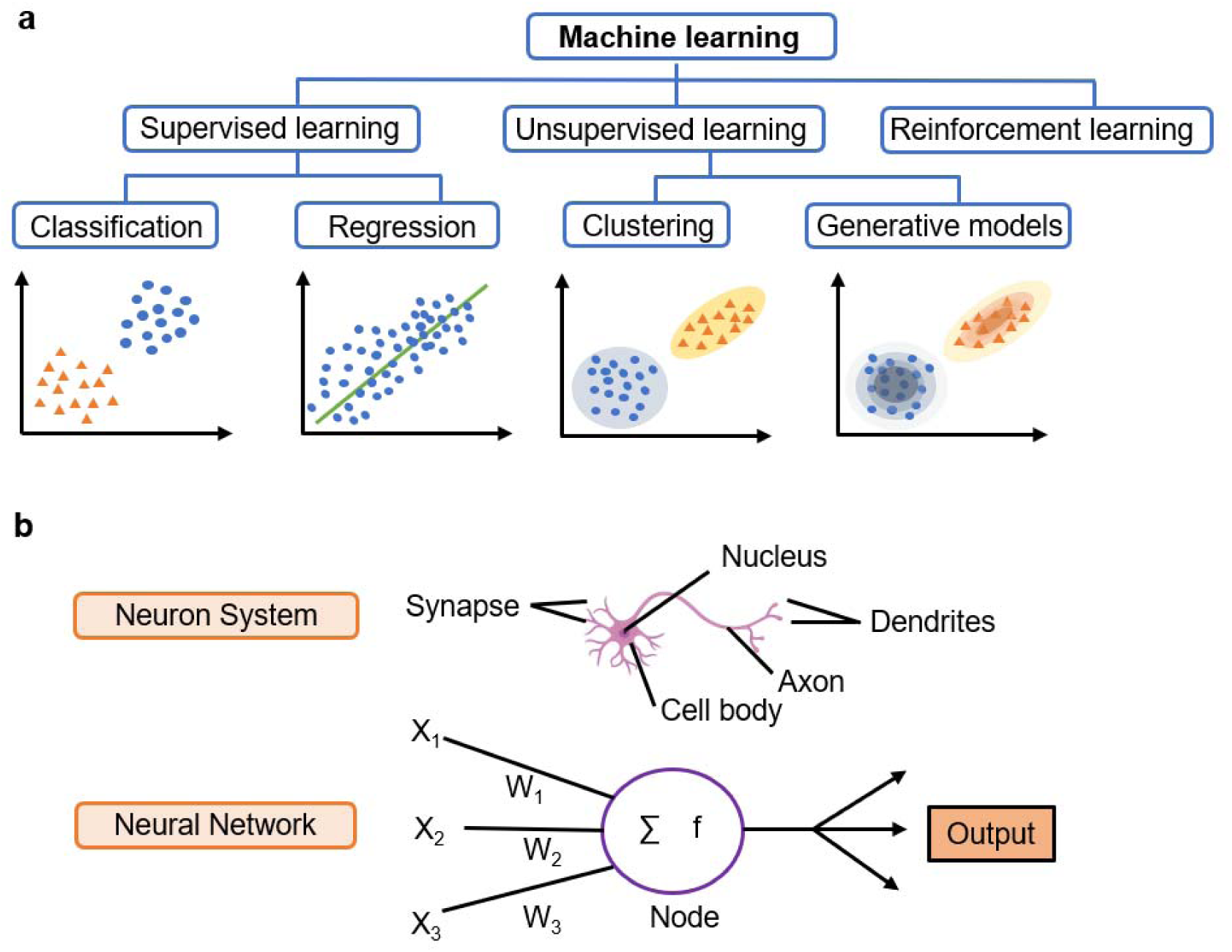
Overview of machine learning and deep learning types and structures. **a** Machine Learning encompasses supervised learning (e.g., regression, classification), unsupervised learning (e.g., clustering, generative models), and reinforcement learning. **b** Structure of neural networks mimicking the human neuron system.

## Results

### Mechanical and physiochemical properties investigation of GelMA-Alginate hydrogel

Achieving a successful bioprinted scaffold relies on biomaterials with optimized rheological properties. Rheological assessments were carried out on various combinations of biomaterials to culture cells encapsulated in hydrogel for extended periods. These hydrogel precursors were tested for rheological characterization to measure crosslinking kinetics and static viscosity. **Fig. 2a** illustrates the rheological behavior of various GelMA-Alginate compositions during the photocrosslinking process. The 5% GelMA formulation exhibited the highest storage modulus upon light exposure, indicating superior elastic properties and network formation. This suggests that the 5% GelMA composition achieves the most efficient crosslinking, resulting in a more robust hydrogel structure.

Samples with a high concentration of methacryloyl groups demonstrated rapid initiation of the photo-crosslinking reaction, as evidenced by the immediate increase. This behavior is characteristic of highly functionalized GelMA, where the abundance of crosslinkable groups facilitates rapid network formation. As the alginate concentration increased, the onset of crosslinking was delayed. However, the storage modulus exhibited a higher value, corresponding to the increased viscosity. This phenomenon can be attributed to the interference of alginate with the GelMA photocrosslinking process. Alginate molecules may physically impede the interaction between methacryloyl groups, resulting in a less densely crosslinked network and, consequently, lower elastic properties.

**Fig. 2:**
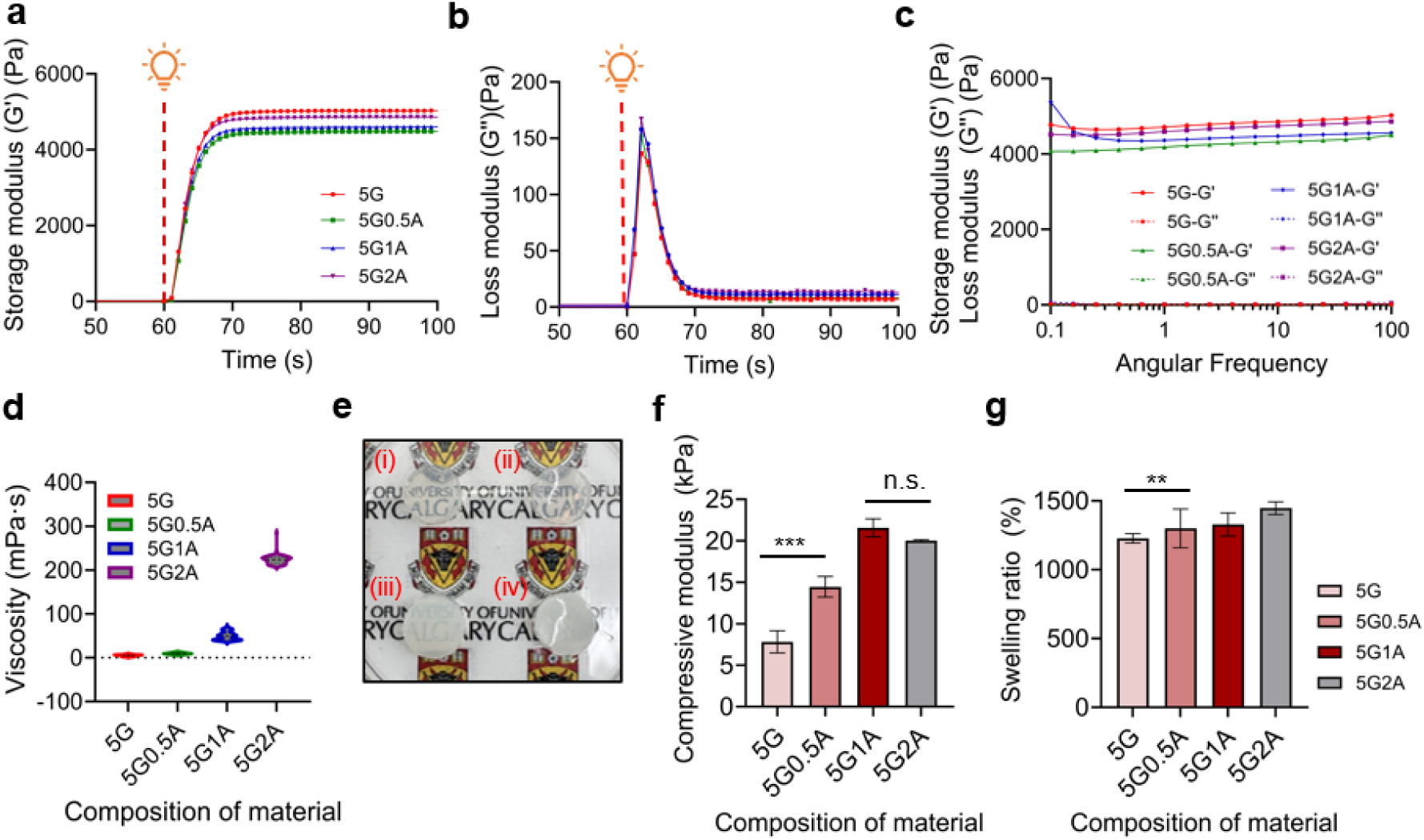
Rheological, mechanical, and physical characterization of GelMA-Alginate hydrogels. **a** Storage modulus at which light exposure begins after 60 seconds. **b** Loss modulus at which light exposure begins after 60 seconds. **c** Angular frequency response to evaluate the crosslinking stability of materials. **d** Static viscosity of materials used in bioprinting for bioprinting parameter input. **e** Optical transmittance of specimens subjected to photocrosslinking under 405 nm wavelength irradiation. **f** Compressive modulus for four distinct material formulations. (n.s. not significant, *** *p* < 0.001). **g** Swelling ratio for four distinct material formulations (** *p* < 0.01).

As the alginate concentration increased, the loss modulus might have slightly increased (**Fig. 2b**). This would be consistent with the decrease in storage modulus, as the alginate interferes with the photocuring process. 5G2A, with its higher loss modulus, indicates that the material exhibits greater viscosity and reduced elasticity compared to 5G. The rate of decrease in loss modulus during photocuring can be slower for samples with higher alginate concentrations. This would reflect the interference of alginate with the crosslinking process, resulting in a more gradual transition from viscous to elastic behavior. The initial loss modulus (before initiating the light exposure) might be higher for samples with higher alginate concentrations due to the increased viscosity contributed by the alginate polymer. These findings highlight the complex interplay between GelMA concentration, methacryloyl group density, and the presence of secondary polymers like alginate in determining the final mechanical properties of photocrosslinked hydrogels. Such insights are crucial for tailoring bioink formulations to achieve the desired mechanical characteristics for specific bioprinting applications.

As shown in **Fig. 2c**, the storage modulus remains relatively constant across a range of angular frequencies, which indicates that the material has achieved a stable and elastic network. All materials used in the test here are fully crosslinked and show a plateau in storage modulus values, showing that the deformation rate does not significantly affect the material’s elastic properties. In addition, for a fully crosslinked material, the loss modulus is significantly lower than the storage modulus across the same range of angular frequencies. This indicates that the material behaves more like a solid (elastic) than a liquid (viscous). The degree of crosslinking in GelMA hydrogels can be effectively assessed by analyzing angular frequency trends, the immediate response to light exposure, and the relative stability of the storage modulus and loss modulus.

Additionally, the static viscosity of the material, one of the critical bioprinting parameters, was investigated. During this measurement, a fluid may exhibit a viscosity plateau around lower values and then display shear thinning as the shear rate increases in a flow curve. Alginate, a high molecular weight polysaccharide, when added to a GelMA solution, increases the average molecular weight of the entire hydrogel precursor. This results in greater entanglement between polymer chains, which is a major factor contributing to the increased viscosity of the solution. As illustrated in **Fig. 2d**, the static viscosity of the GelMA-based hydrogel precursor increases with higher alginate concentration. Such high viscosity positively impacts printing fidelity and the ability to maintain the structural integrity of the printed constructs ^30,31^.

To evaluate the transparency, the logo remained clearly discernible in the 5G and 5G0.5A samples (**Fig. 2e**). Conversely, the 5G1A sample exhibited reduced logo visibility, while the 5G2A sample demonstrated complete opacity, fully obscuring the underlying logo. These observations indicate a progressive increase in opacity correlating with the compositional variations among the samples, with a notable rise in opacity as the alginate ratio is increased.

As the alginate concentration increases, the initial mechanical strength of the hydrogel improves. However, excessive alginate concentrations can interfere with the photo-crosslinking of GelMA. **Fig. 2f** illustrates that the mechanical strength increases up to the 5G1A composition but slightly decreases for 5G2A. This suggests that alginate may impede light transmission during the photocrosslinking process or interfere with the interaction of GelMA’s methacrylate groups.

Due to alginate’s high hydrophilicity, its incorporation into GelMA enhances the porosity of the hydrogel network, generally leading to an increased swelling ratio ^32,33^. As illustrated in **Fig. 2g**, the swelling ratio increases with the addition of alginate to the 5G formulation.

### Development and validation of the modified bioprinting platform

A conversion of a commercially available 3D printer into a bioprinter is detailed in Shin *et al* ^12^. It was a modified version of a low-cost 3-axis fused deposition modeling printer (Ender 3 Pro, Creality 3D, China) integrated with a pneumatic-based liquid dispenser (FEITA 983, FEITA, China). The enhanced setup includes a fast-switching circuit (**Supplementary Fig. S1a**), a programmable linear stage (**Supplementary Fig. S1b**), LED lighting and cell mixer (**Supplementary Fig. S1c**), and a modified liquid dispenser (**Supplementary Fig. S1d**).

To accommodate the data-intensive nature of ML training, a movable printing stage and additional modifications were incorporated into the original 3D bioprinting platform (**Fig. 3a**). The G-code was programmed to utilize the bed stepper motor, shifting the printing linear stage rightward after each droplet deposition, thereby facilitating the acquisition of new cell-laden droplet image data (**Supplementary Movie S1 and Fig. S2**). A USB digital camera (Jiusion, Shenzhen, China) was integrated and connected to the laptop to capture droplets’ images to enable real-time monitoring and analysis. This integrated system allows for detailed visualization of the droplets (**Supplementary Movie S2**). The syringe was connected to a liquid dispenser, and the setup employed a modified circuit board to ensure precise control over the air pressure output. This pneumatic extrusion-based method produces significantly smaller droplets (0.1 μL) compared to piston, screw, or syringe pump-based methods (**Supplementary Movie S3**). Overall, it has demonstrated precise control over various droplet sizes with high repeatability. This configuration ensures that the developed bioprinting system consistently maintains the predetermined air pressure settings throughout the entire bioprinting process. For consistent cell number bioprinting, we developed and manufactured a motorized button capable of inserting a 10 mm x 1.5 mm stir bar into the syringe to mix the bioink homogenously (**Supplementary Movie S4**). In the video, PBS mixed with glitter was used instead of cells in the syringe for visualization purposes.

**Fig. 3:**
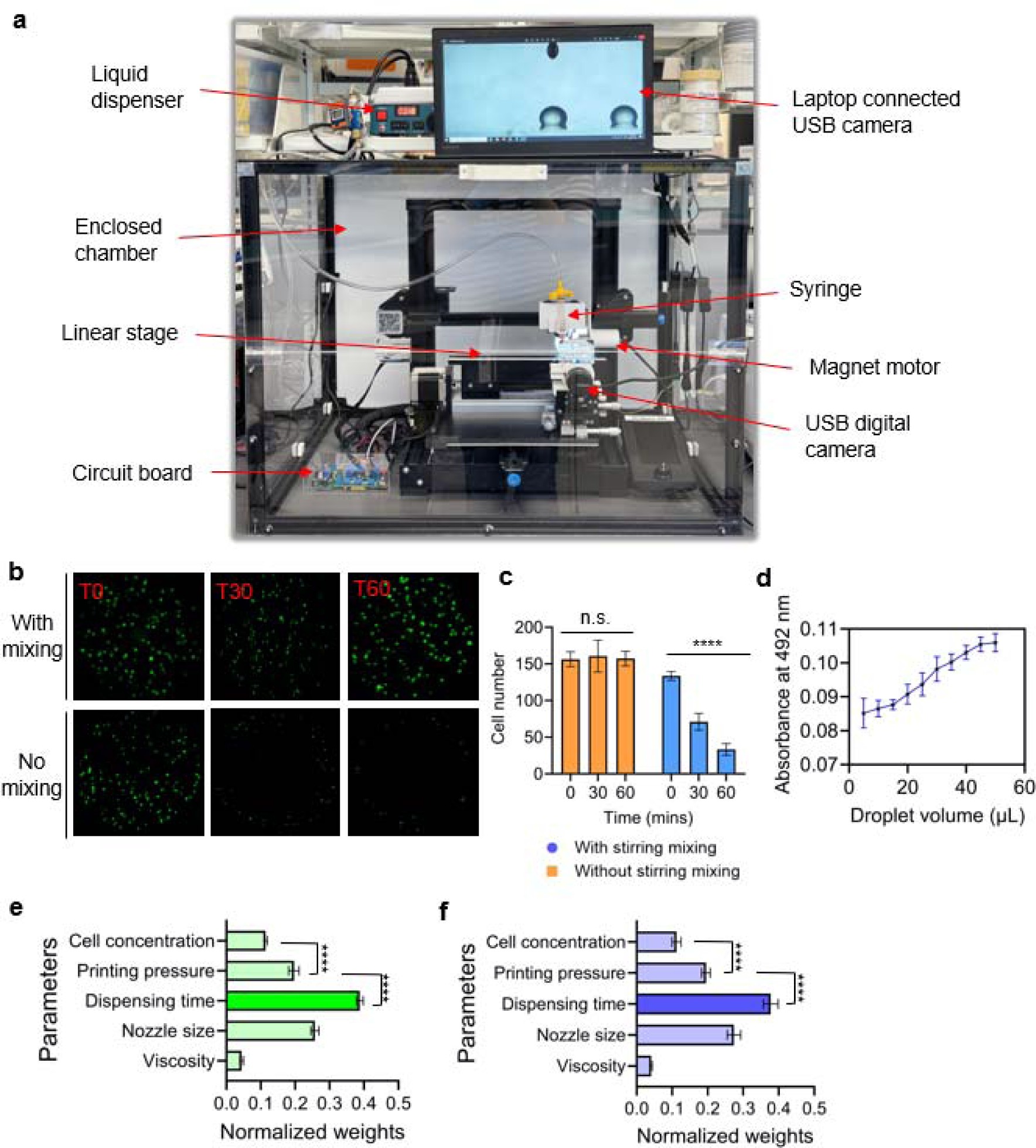
**a** Modified 3D bioprinting setup featuring. **b** Impact of in-syringe bioink homogenization on cell distribution and viability in bioprinted constructs. **c** Comparison of bioprinted cell numbers over time frames (0, 30, 60 minutes) to assess the safety of cell numbers with and without the stirring magnet (n.s.: not significant, **** *p* < 0.0001). **d** Absorbance results at 492 nm based on cell number resulting from droplet volume variations. **e** Feature weight of decision tree algorithm (**** *p* < 0.0001). **f** Feature weight of random forest algorithm (**** *p* < 0.0001).

In **Fig. 3b**, the dispensed bioink was immediately crosslinked and imaged under a microscope at three time points: at the start of printing (T0), 30 minutes later (T30), and 60 minutes later (T60) (**Supplementary Movie S5**). This result illustrates that cells with the mixing system maintained a constant cell number over time, whereas cells dispensed without the mixing system showed a decrease in cell number within the droplets over time. To comprehensively analyze the effects of two independent variables (the presence of a magnetic stirring system and printing duration) on a single dependent variable (cell number), we employed a Two-way ANOVA. This statistical method allows for the simultaneous examination of the main effects of each independent variable and their potential interaction effect on the cell number. **Fig. 3c** illustrates the variation in the number of printed cells depending on whether the syringe used for printing is equipped with a magnetic stirring system. With the stirring system, there was no statistically significant difference between the initial cell count at the start of printing and the cell number in the printed cellular droplets even after 60 minutes of continuous printing. Conversely, in the absence of the stirring system, a statistically significant decrease was observed in the cell number of the cellular droplets printed after 60 minutes compared to the initial cell number. Ultimately, a stirring system resulted in a statistically significant difference in cell number.

The printed GFP-tagged 3T3 cell-laden droplets were produced in various sizes (up to 50 μL) on a 96-well plate. The fluorescence signal was measured at an absorbance of 492 nm using a plate reader to quantify the intensity of fluorescence emitted by samples containing the GFP-tagged cells. By measuring the fluorescence intensity, the plate reader provides quantitative data on the number of cells present in each droplet. The results demonstrated that as the droplet size increased, the fluorescence signal also increased, corresponding to a higher number of cells (**Fig. 3d**). This indicates that the developed bioprinting system not only produced uniform and stable droplets but also automated the data collection process, allowing for high-throughput analysis of multiple samples at the same time. This efficiency is particularly beneficial in experiments requiring large datasets for ML training.

Additionally, this study focused on identifying and quantifying the relative importance of bioprinting parameters in determining the final volume of bioprinted droplets (**Table 1**). Using a systematic approach, each bioprinting parameter varied independently while keeping others constant to assess its effect on droplet volume. All parameters have a statistically significant effect on droplet volume (**Supplementary Table S2**). **Fig. 3e, f** shows the weight of each bioprinting parameter. As a result, dispensing time emerged as the most influential factor in determining droplet volume among the parameters. The nozzle size and printing pressure were the next most significant parameters. Under the assumption that the primary parameters are precisely controlled, the study revealed that material viscosity and cell concentration exert comparatively minor effects on droplet volume regulation. These findings provide valuable insights into the hierarchical importance of bioprinting parameters, potentially enabling more precise control over droplet volume in bioprinting processes. Such knowledge is crucial for optimizing bioprinting protocols and achieving higher levels of precision in fabricating complex tissue constructs.

**Table 1.** Bioprinting parameters include viscosity of biomaterial, nozzle size, printing time, printing pressure, and cell concentration to acquire the desired droplet volume.

| Independent variables |  |  |  |  | Dependent variables |
| --- | --- | --- | --- | --- | --- |
| Viscosity (mPa·s) | Nozzle gauge (I.D. mm) | Printing time (s) | Printing pressure (psi) | Cell concentration | Droplet volume (μL) |
| 10.03 | 23 (0.337) | 0.05 | 1.5 | 2.8×10<sup>6</sup>/3 mL | ... |
| 24.66 | 25 (0.26) | 0.1 | 2 | 5.6×10<sup>6</sup>/3 mL |  |
|  | 27 (0.21) | 0.15 |  |  |  |

### Image processing and droplet volume calculation

To achieve uniform volume of droplets during bioprinting and maintain consistent printing conditions, the glass slide substrate was treated with a hydrophobic saline coating (**Fig. 4a**). This treatment allowed for the generation of high-throughput and uniform droplets on a standard microscopy glass slide. Specifically, 1H, 1H, 2H, 2H-Perfluorooctyltrichlorosilane (Fisher Scientific, Waltham, MA, USA) was used to coat the surface of the glass slides, resulting in hydrophobic surfaces. The glass slides were then placed on a movable linear stage, enabling droplet formation. Then, cell-laden bioinks (5G1A and 5G0.5A) were loaded in a 3 mL syringe equipped with a small magnetic stir bar to ensure homogeneous cell distribution in the bioink throughout the droplet printing process (**Fig. 4b**).

**Fig. 4:**
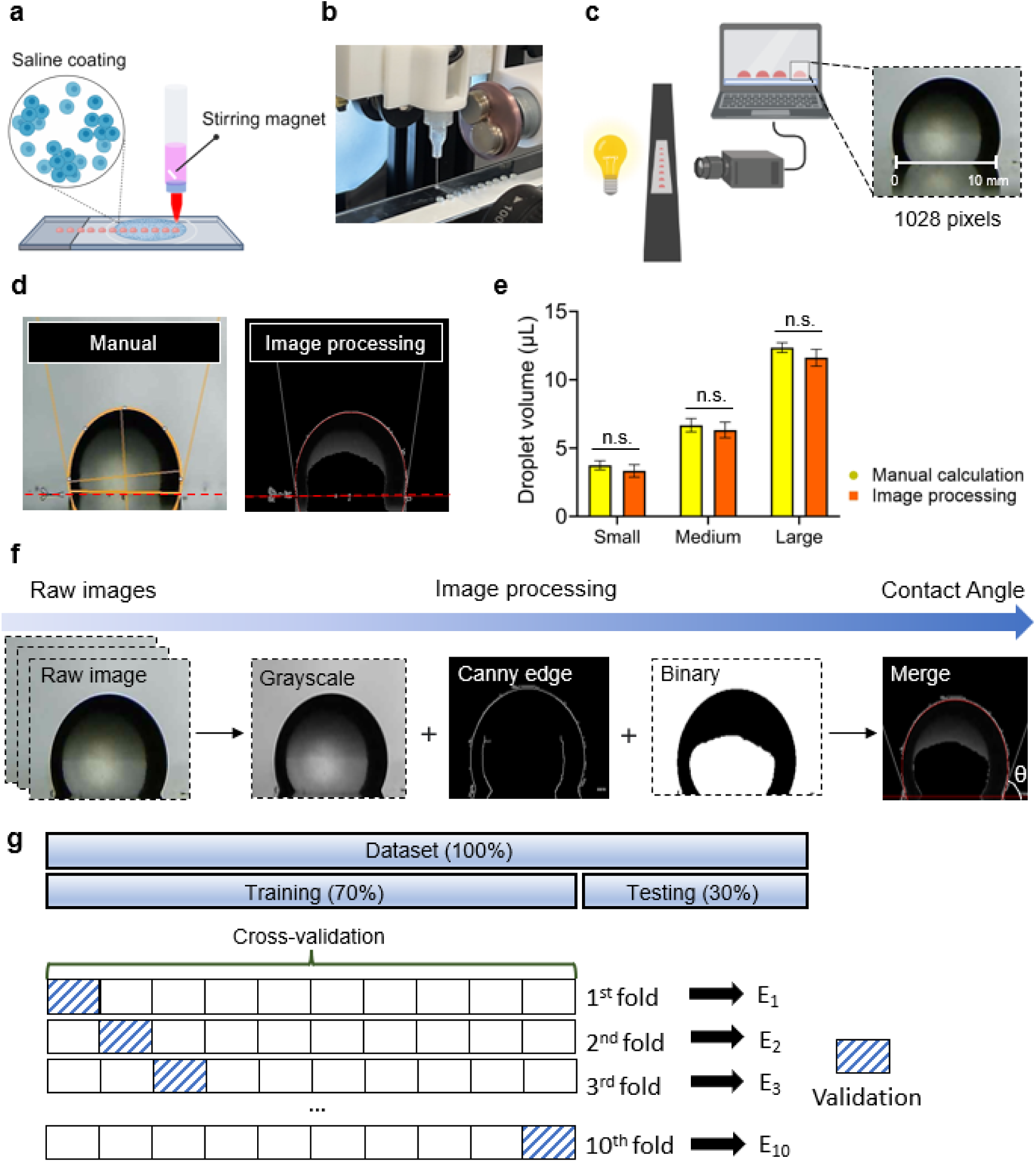
**a** Bioprinting multiple cellular droplets in a high-throughput manner during a single bioprinting run on a saline-coated glass slide. **b** Demonstration of consistency within bioprinted GFP-tagged 3T3 fibroblast cellular droplets. **c** Illustration depicting the calibration method and process for measuring droplet volume in a droplet volume measurement system. **d** Comparative analysis of manual and automated droplet size measurements across three different size categories (n.s.: not significant). **e** Comparison between droplet volumes calculated manually using ImageJ and those measured using the developed image processing method (*n*=10). **f** Overall image preprocessing process for preparing the dataset for machine learning training. **g** Data partitioning and validation strategy: An initial 70:30 split of the dataset into training and testing sets, followed by the implementation of 10-fold cross-validation on the training set to mitigate overfitting.

A microruler was fixed at the droplet dispensing location to accurately measure the droplets’ actual length at the dispensing point, yielding a value of 10 mm. The corresponding pixel count of this length in the images was found to be 1028 pixels (**Fig 4c**). Finally, this calibration resulted in a conversion factor of 0.009728 mm per pixel, enabling accurate translation of pixel measurements to actual dimensions.

To demonstrate the robustness of our droplet size calculation software, we conducted a comparative analysis with the traditionally widely used droplet volume measurement calculator in ImageJ software (**Fig. 4d**). This analysis utilized measurements obtained from images of 10 randomly selected droplets. The analysis encompassed droplets of various sizes: small droplets with a mean volume of 3.54 μL, medium droplets averaging 6.54 μL, and large droplets with a mean volume of 11.98 μL. As a result of the comparative analysis, there was no statistically significant difference between the droplet volume measurements obtained using the ImageJ software method and those derived from the image processing method developed in Python in this study (**Fig. 4e**).

Image processing employs a series of techniques to enhance efficiency and accuracy. Precise edge detection was required to enhance droplet volume calculation accuracy. Raw images underwent three extraction techniques (e.g., grayscale, Canny edge, and binary method), combined to produce the final image (**Fig. 4f**). Initially, grayscale conversion was applied to reduce computational complexity by transforming color images into single-channel representations. Then, Canny edge detection was utilized to identify image boundaries and obtain clear outlines of the objects. Lastly, binary thresholding was implemented to simplify the image to black and white pixels, effectively reducing noise and irrelevant details. By employing these techniques sequentially, the image processing pipeline achieved both simplification of color images and enhanced processing efficiency, providing a robust foundation for subsequent analysis tasks. The diameter and contact angles were measured from the final image, and the droplet volume was calculated using the following **Equation (1)**:

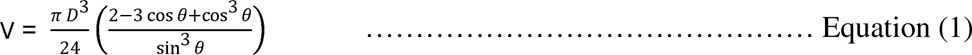

By combining these processed images, we achieved reliable and consistent volume measurements across all samples, demonstrating the robustness of the proposed approach in capturing fine details necessary for precise droplet quantification. The droplet volumes obtained were extracted and transferred to Excel, and the data was partitioned into training and testing to train three ML and two DL algorithms (**Supplementary Fig. S3**). The total 1758 image datasets were divided into a 70% training set and a 30% testing set (**Fig. 4g**). To mitigate overfitting, 10-fold cross-validation was performed to ensure the performance estimates are robust and that the models will generalize properly to new droplet images. The dataset was randomly divided into 10 equal-sized subsets or folds, and then the model was trained on 9 folds and validated on the remaining 1-fold. This process was repeated 10 times, with each fold as the test set once. Finally, the model’s performance was averaged across all 10 iterations.

### Optimization of the hyperparameters

Optimal hyperparameter tuning is essential for maximizing the performance of ML models. By leveraging various optimization methods, such as grid search, random search, bayesian optimization, gradient-based methods, evolutionary algorithms, and hyperband, practitioners can efficiently navigate the hyperparameter options and identify the best configurations for their specific use cases.

To evaluate the performance of the algorithms in predicting printing parameters, our study was structured into two distinct phases: Phase 1 and Phase 2. During Phase 1, an optimization process was conducted to determine the optimal hyperparameters for each algorithm. This involved utilizing 10-fold cross-validation to identify the hyperparameters that yielded the best performance. Utilizing k-fold cross-validation can mitigate the bias and variance associated with a single data split, leading to a more robust evaluation of model performance ^34^. Additionally, it aids in detecting overfitting issues and facilitates the discovery of a generalized model, with each algorithm exhibiting distinct optimized hyperparameters. In **Fig 5a**, the DT performed optimally with a maximum depth of 7, while the RF demonstrated superior performance with 10 estimators (**Fig. 5b**). Notably, given the multitude of hyperparameters in RF, a systematic approach employing GridSearchCV, an automated hyperparameter detection function, was utilized to determine the most optimal combination. Among the identified combinations, the parameter with the highest weighting, namely the number of estimators, was selected as the representative hyperparameter. Deeper architectures can contribute to more accurate classification; however, this intensifies the risk of overfitting, necessitating the trimming of questions to an appropriate level. PR yielded optimal results with a degree of 7 (**Fig. 5c**), MLP with a maximum epoch of 100 (**Fig. 5d**), and LSTM achieved the peak performance with a maximum epoch of 100 (**Fig. 5e**).

**Fig. 5:**
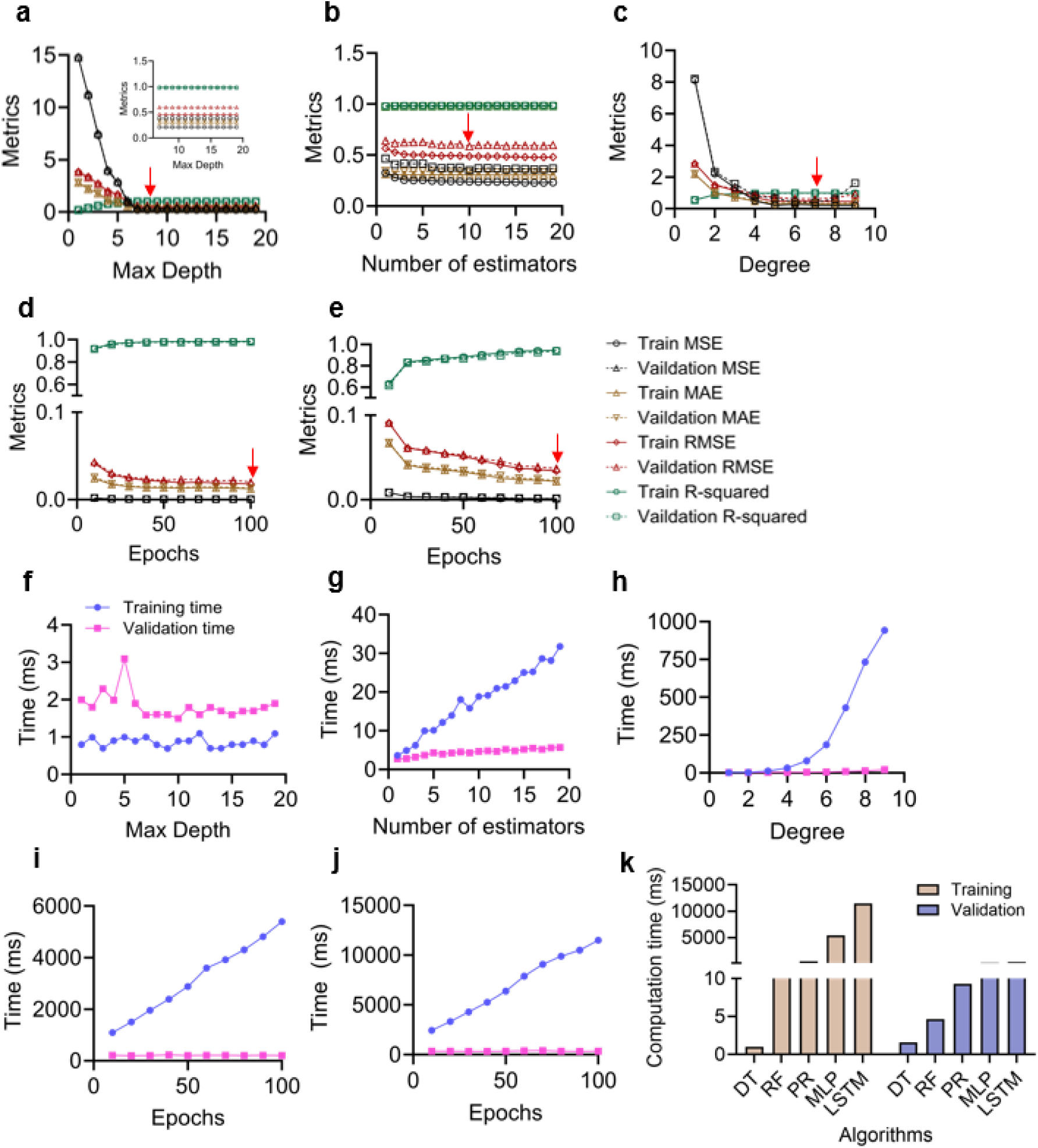
**a** Optimization process to find the optimized hyperparameters for a decision tree. **b** Process to find the optimized hyperparameters for a random forest. **c** Process to find the optimized hyperparameters for a polynomial regression. **d** Process to find the optimized hyperparameters for a multi-layer perception. **e** Process to find the optimized hyperparameters for a long short-term memory. **f** Training and validation time when optimizing the hyperparameters in the decision tree. **g** Training and validation time for the random forest. **h** Training and validation time for polynomial regression. **i** Training and validation time for multi-layer perception. **j** Training and validation time for long short-term memory. **k** Comparison of computation time for all algorithms at the moment of optimized hyperparameter settings (*n*=1).

### Computation time for hyperparameter optimization

Hyperparameter optimization often involves exploring many combinations of hyperparameters. The time required to evaluate each combination directly impacts the overall optimization process. Typically, training time is longer than validation time due to the iterative nature of the training process and the complexity of the models involved (**Fig. 5f-j**). Contrary to typical expectations, the DT model exhibited longer validation than training times. The validation phase, especially with k-fold cross-validation, does not benefit from this same level of parallel processing. As a result, the validation phase can take longer than the training phase. However, DT algorithms train quickly because they can build multiple trees at the same time. It achieved hyperparameter optimization with the fastest computation time compared to other algorithms when optimizing the hyperparameters (**Fig. 5k and Table 2**).

**Table 2.** Computation times of three machine learning and two deep learning algorithms using the dataset at the moment of optimized hyperparameter. Training and validation time for the 10-cross-validation process.

| Algorithms | Training time (ms) | Validation time (ms) |
| --- | --- | --- |
| Decision Tree | 0.998282 | 1.591182 |
| Random Forest | 18.855333 | 4.672742 |
| Polynomial Regression | 430.593181 | 9.286737 |
| Multi-layer perception | 5394.694638 | 205.264473 |
| Long short-term memory | 11486.32846 | 335.380983 |

The complexity of the model, such as the depth of a neural network or the number of trees in an RF, directly impacts the duration of the training. More complex models require more time to converge to an optimal solution. Therefore, the computation time of deeper models such as MLP and LSTM takes longer than ML algorithms (e.g., DT, RF, and PR) (**Fig. 5k**). In summary, while both training and validation are essential components of hyperparameter optimization, training typically demands more time due to its iterative nature and the need for parameter updates. Conversely, validation is faster as it evaluates the model’s performance without further adjustments. Efficient optimization techniques and proper resource management can help balance the time requirements of these phases, ultimately leading to more effective and timely model development.

### Comparative evaluation of machine learning algorithms for droplet volume prediction

The hyperparameters identified as optimal during the initial optimization phase were applied to each algorithm. Subsequently, each model underwent a rigorous training and evaluation process of 10 independent iterations. To compare the performance of the algorithms, we evaluated three ML and two DL algorithms using the following metrics: MAE, MSE, RMSE, and R-squared (**Table 3**). First, MAE is a useful indicator for analyzing the model’s prediction accuracy and error magnitude. As shown in **Fig. 6a**, there was no statistically significant difference between the ML algorithms (DT, RF, and PR) and no significant difference between the DL algorithms (MLP and LSTM). However, a statistically significant difference was observed between the ML and DL algorithms, with a p-value of 0.0000108. This shows that the MLP and LSTM models have a smaller MAE than the DT, RF, and PR models, meaning their predictions are closer to the actual values. As a result, MLP and LSTM demonstrate better prediction accuracy and overall performance.

**Fig. 6:**
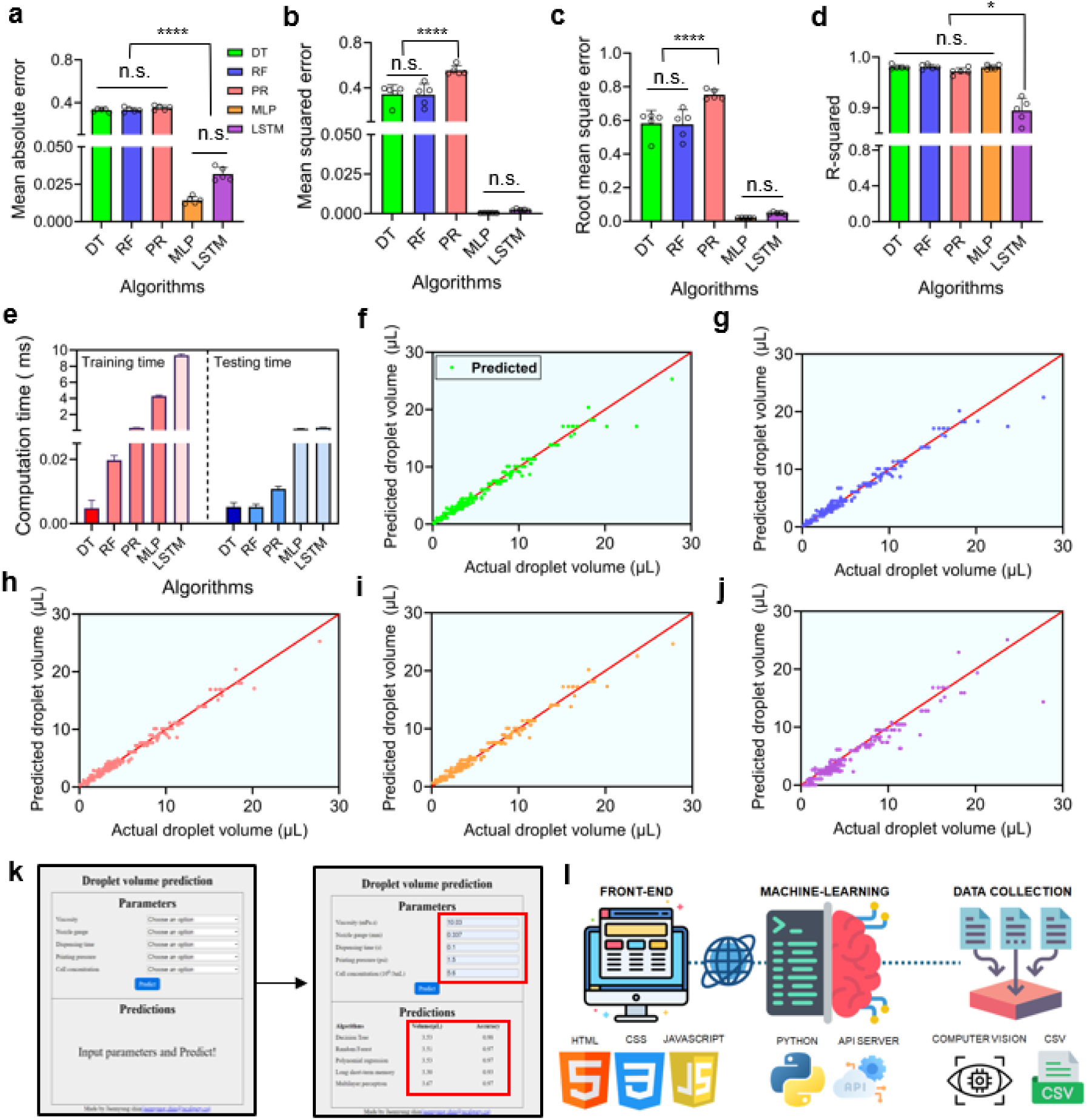
Performance evaluation of machine learning and deep learning models. **a** Mean absolute error across models (n.s.: not significant, **** p < 0.0001). **b** Mean squared error comparison (n.s.: not significant, **** p < 0.0001). **c** Root mean square error analysis (n.s.: not significant, **** p < 0.0001). **d** R-squared values (n.s.: not significant, * *p* < 0.05). **e** Computational time for training and testing phases (*n*=5). **f** Visualization of model performance by comparing the actual and predicted values of the decision. **g** Random forest: comparison of actual and predicted values. **h** Polynomial regression: comparison of actual and predicted values. **i** Multi-layer perception: comparison of actual and predicted values. **j** Long short-term memory: comparison of actual and predicted values. **k** Implementation of a web-based interface for the droplet volume measurement system, optimized using machine learning and deep learning algorithms. **l**. Schematic representation of the full-stack web application architecture.

**Table 3.**
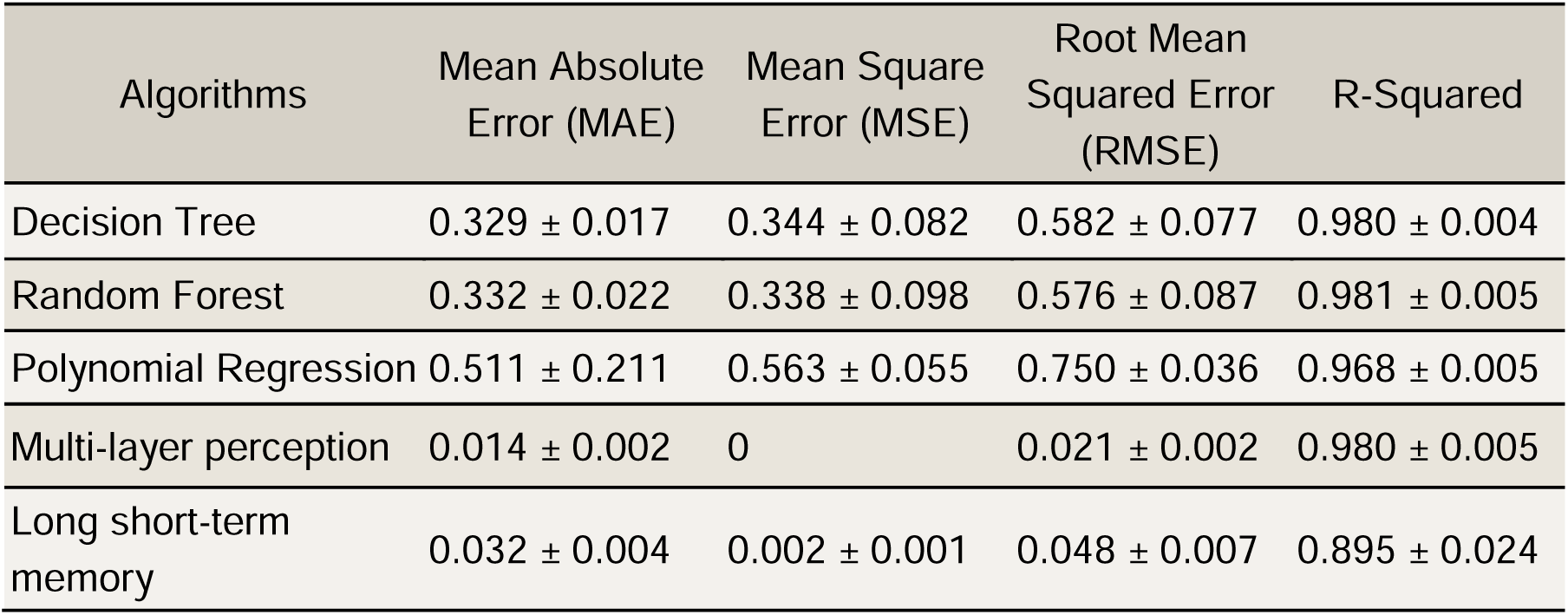
Performance evaluation of three machine learning and two deep learning algorithms.

| Algorithms | Mean Absolute Error (MAE) | Mean Square Error (MSE) | Root Mean Squared Error (RMSE) | R-Squared |
| --- | --- | --- | --- | --- |
| Decision Tree | $0.329 \pm 0.017$ | $0.344 \pm 0.082$ | $0.582 \pm 0.077$ | $0.980 \pm 0.004$ |
| Random Forest | $0.332 \pm 0.022$ | $0.338 \pm 0.098$ | $0.576 \pm 0.087$ | $0.981 \pm 0.005$ |
| Polynomial Regression | $0.511 \pm 0.211$ | $0.563 \pm 0.055$ | $0.750 \pm 0.036$ | $0.968 \pm 0.005$ |
| Multi-layer perception | $0.014 \pm 0.002$ | 0 | $0.021 \pm 0.002$ | $0.980 \pm 0.005$ |
| Long short-term memory | $0.032 \pm 0.004$ | $0.002 \pm 0.001$ | $0.048 \pm 0.007$ | $0.895 \pm 0.024$ |

Secondly, the MSE is calculated by averaging the squared differences between predicted and actual values. Lower MSE values indicate closer alignment between model predictions and actual values. The DL models, MLP and LSTM, demonstrated the lowest MSE values with no statistically significant difference, indicating superior performance **Fig. 6b**. MSE is particularly sensitive to outliers because it uses squared differences in calculations. The PR model, which shows relatively higher values, may have captured outliers less effectively than other algorithms. However, it is noteworthy that all models in this study achieved MSE values below 1, suggesting generally good performance across the board ^35^. It is important to consider that while MSE provides valuable insights into model performance, it should be interpreted in conjunction with other metrics to evaluate model efficacy comprehensively.

Thirdly, RMSE mitigates the distortion caused by squaring the errors in MSE by taking the square root, making it more interpretable. RMSE is more robust against outliers compared to MAE. As shown in **Fig. 6c**, the DL models exhibit the lowest RMSE values, indicating superior performance. For ML models, there is no statistically significant difference between the DT and RF models. However, a statistically significant difference was observed between the PR models. Notably, all models evaluated in this study achieved RMSE values below 1.

Lastly, R-squared is an effective evaluation metric for comparing relative performance, as it represents the proportion of variance in the actual values explained by the predicted values. The closer the R-squared value is to 1, the better the model’s performance; conversely, a value closer to 0 indicates a less accurate model ^35^. As shown in **Fig. 6d**, the models DT, RF, PR, and MLP all exhibit R-squared values close to 1 (0.980, 0.981, 0.968, and 0.980, respectively), with no statistically significant differences among them, indicating excellent performance. However, the LSTM model shows a slightly lower R-squared value of 0.895. Despite this, it is still close to 1, suggesting that all the trained models demonstrate sufficient performance overall.

### Computation times required for the machine learning algorithms

Optimization of computation time is an essential factor that directly affects the model’s performance and the efficiency of the development process, as well as cost-effectiveness and practical applicability. DL models often require longer training times to fully leverage the depth and complexity of neural networks compared to ML models (**Fig. 6e and Table 4**). This can lead to significant performance improvements, especially for tasks that require high-level feature extraction. As shown in **Fig. 6e**, the MLP model took 4.296 seconds for training and 0.218 seconds for testing, while the LSTM model required 9.349 seconds for training and 0.350 seconds for testing. Longer training times allow the model to converge more effectively, reducing the loss function and improving accuracy. However, excessively long training times can lead to overfitting, where the model performs well on the training data but poorly on unseen data. These risks can be mitigated through regularization techniques and early stopping ^36^.

**Table 4.**
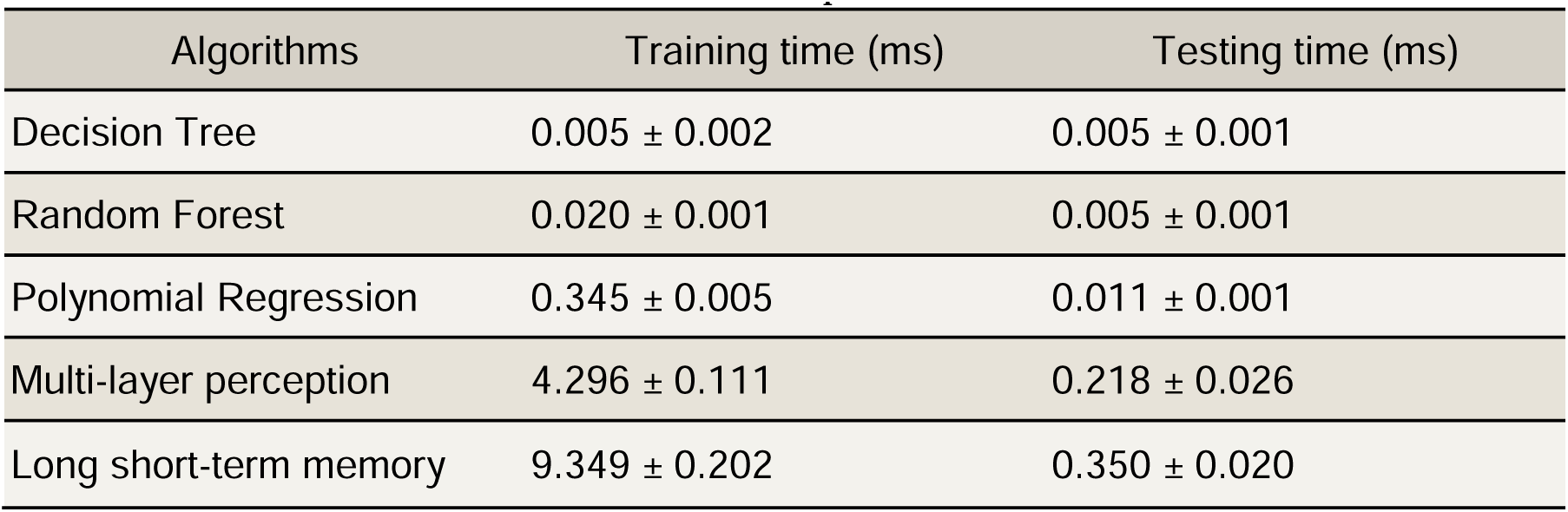
Computation times of three machine learning and two deep learning algorithms were measured on the dataset to evaluate their performances.

Additionally, computation time is directly related to cost. While longer training times can improve performance, they also increase computational expenses. It is crucial to balance training duration with available resources for practical applications. Ultimately, increased training time can enhance model performance by enabling better convergence, thorough hyperparameter optimization, and robust validation. However, balancing this with the risk of overfitting and real-world constraints on computational resources is essential ^36^.

### Development of user-interface web application

Building on the algorithm’s demonstrated high prediction accuracy, the predicted droplet volumes are in close alignment with the actual measured volumes, indicating the model’s effectiveness in accurately capturing the characteristics of the droplets. This consistency between predicted and actual values underscores the algorithm’s reliability in practical applications (**Fig. 6f-j**). In addition, we developed a front-end web application based on these models, as illustrated in **Fig. 6k**. This user-friendly interface leverages the high-performance predictive model to provide real-time, accurate estimations of droplet volumes. The web interface is divided into two main sections: Parameters and Predictions. The parameters section includes viscosity, nozzle size, dispensing time, printing pressure, and cell concentration. By adjusting these parameters, users can obtain predicted droplet volumes before cellular droplet bioprinting, facilitating a faster, easier, and more precise acquisition of the desired cellular droplets. The prediction section presents results in three columns: algorithm, volume, and accuracy, with predictions generated using five different algorithms. The algorithm column specifies the type of algorithm used, the volume column shows the predicted droplet volume, and the accuracy column indicates the precision of each algorithm. Data for droplet size prediction was gathered using computer vision techniques and equations, with the results compiled in CSV format (**Fig. 6l**). This CSV data was then used as the training dataset for the ML algorithms, which were implemented using Python.

To provide access to the trained ML algorithms as a web service, we developed an application programming interface (API) server. The web application communicates with this API server, enabling users to interact with the prediction models via a user-friendly web interface. Through this interface, users can input parameters, submit requests to the server, and receive predictions for droplet sizes. This integrated system allows researchers to efficiently predict and optimize cellular droplet volumes prior to bioprinting, significantly enhancing the precision and effectiveness of the bioprinting process.

### Assessment of cellular viability and proliferative capacity in 3D bioprinted constructs

Maintaining cell viability while minimizing shear stress-induced damage at the nozzle is critical. As illustrated in **Fig. 7a**, GFP-3T3 cells encapsulated in 5G and 5G0.5A hydrogels exhibited numerous cell elongations, indicating favorable cell-matrix interactions. Furthermore, proliferation assays conducted up to day 7 revealed a steady increase in cell numbers (**Fig. 7b**). In contrast, cells in the 5G material initially showed higher proliferation rates, and by day 3, both 5G and 5G0.5A compositions supported comparable proliferation kinetics. Cellular morphology and nuclear integrity were further confirmed through phalloidin and dapi staining, respectively (**Fig. 7c**). Quantitative analysis of cell viability from day 1 to day 5 consistently demonstrated viability rates exceeding 90% (**Fig. 7d**).

**Fig. 7:**
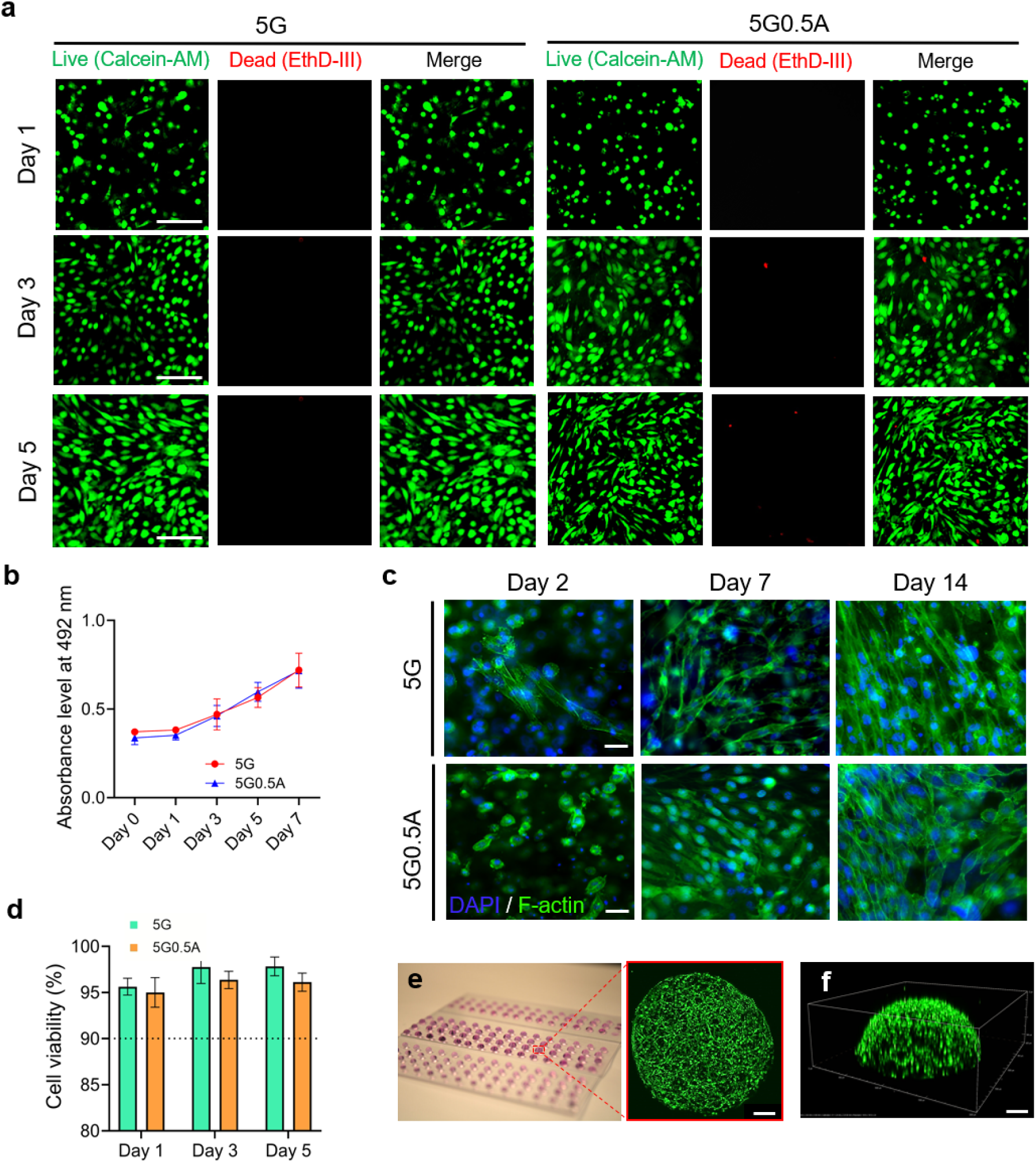
**a** Representative images showing live (green) and dead (red) NIH 3T3 fibroblast cells at days 1, 3, and 5. Scale bars = 100 µm. **b** Cell proliferation rate from day 0 to day 7. **c** 3T3 cells embedded in GelMA-Alginate hydrogels were observed over two weeks under conditions 5G and 5G0.5A. Cell nuclei were stained with DAPI (blue) to represent the original cells, while the cytoskeleton was visualized using phalloidin (green). Scale bars = 50 µm. **d** Cell viability, determined by counting live and dead cell populations on days 1, 3, and 5. **e** Maximum intensity projection images through 1 mm thickness of bioprinted GFP-3T3 cell encapsulated in 5G2A bioink. Scale bar = 200 um. **f** 3D confocal imaging of GFP-3T3 cells encapsulated in 5G2A bioink. Scale bar = 200 um.

To observe the overall distribution of cell growth within the 3D droplet, GFP-3T3 cells were encapsulated in droplets with a thickness ranging from 500 µm to 1 mm. These droplets were then bioprinted and cultured in an appropriate medium. The time-lapse video on day 3 demonstrates the elongation of cells as they proliferate from the bottom to the top area of the droplet (**Supplementary Movie S6**). On day 3, after bioprinting, we performed confocal imaging of GFP-3T3 cells encapsulated in the bioink scaffold. **Fig. 7e** presents maximum intensity projection images of a droplet approximately 1 mm thick, captured at 10X magnification using 9 tiled images. **Fig. 7f** displays a 3D view of the same sample, illustrating the complete elongation of the cells by day 3. These results demonstrate both the hydrogel formulations’ biocompatibility and our bioprinting system’s cell-preserving capabilities.

## Discussion

Droplet bioprinting of cell aggregates and organoids at the microliter scale presents significant challenges ^37^. It is essential to optimize bioprinting parameters before the bioprinting process, maintain precise control during bioprinting, and ensure the high repeatability and stability of the resulting cell-laden constructs ^38^. Therefore, we proposed a strategy for optimizing bioprinting parameters by leveraging ML and DL technologies.

Among bioprinting parameters, bioink composition cannot be overemphasized in extrusion-based bioprinting ^39^. Low concentrations of hydrogel bioink often result in inadequate mechanical stability and printability, complicating the creation of constructs with microliter-sized droplet volumes. Initially, 5G, 5G0.5A, 5G1A, and 5G2A were tested for material characterization. Based on these results, 5G0.5A and 5G1A were identified as the most suitable candidates for droplet bioprinting, and thus were selected for further optimization Samples with a high concentration of methacryloyl groups demonstrated rapid initiation of the photo-crosslinking reaction, as evidenced by the immediate increase in storage modulus. This behavior is characteristic of highly functionalized GelMA, where the abundance of crosslinkable groups facilitates rapid network formation ^40^. Increasing alginate concentration correlated with a slight decrease in the final storage modulus. This phenomenon can be attributed to the interference of alginate with the GelMA photocrosslinking process. Alginate molecules may physically impede the interaction between methacryloyl groups, resulting in a less densely crosslinked network and, consequently, lower elastic properties.

Regarding the 3D bioprinter, we customized it to improve the bioprinting process’s controllability and to collect a high-throughput image dataset for optimization training. This bioprinter enabled the controlled bioprinting of microliter-sized cell-laden droplet constructs. We also installed a cell stirring system within the syringe to ensure a consistent cell number in each droplet. Its effectiveness was verified by checking the cell number at three different time points during the bioprinting process. Additionally, bioprinting was conducted on a hydrophobic glass slide surface to enhance the clarity and repeatability of the droplet image edges. This approach not only increased the contact angles of the droplets to over 90 degrees by making the surface hydrophobic but also facilitated bonding between the hydrogel precursor and the glass, which is suitable for long-term culture and ensures the constructs remain attached for extended periods ^41,42^.

The high-throughput images collected were transferred to a computer for further analysis. Three distinct image processing techniques were applied to these images. By employing multiple image processing methods, we mitigated potential inaccuracies inherent in a single technique, thereby enhancing the robustness of our volume measurements. This comprehensive image analysis streamlines not only provided a high degree of control over the bioprinting process but also contributed to the optimization and standardization of our bioprinting protocols.

The diverse droplet volumes produced by adjusting bioprinting parameters were input into three ML algorithms and two DL algorithms for training and testing. Before this, we performed hyperparameter optimization for each algorithm to identify the configuration that yielded the best predictive performance. By employing hyperparameter optimization, each algorithm was tuned, ensuring the highest predictive accuracy for droplet volume outcomes. In addition, the model’s generalization performance is enhanced, decreasing the risk of overfitting. This rigorous approach allowed us to evaluate and improve our bioprinting process’s performance systematically. The combination of multiple algorithms provided a comprehensive understanding of the parameters influencing cell-laden droplet formation, facilitating the development of robust predictive models. These models are crucial for optimizing bioprinting parameters in real-time, thereby improving the consistency and quality of bioprinted constructs.

The predictive performance was evaluated based on two criteria: droplet volume prediction accuracy and computation time. For the first criterion, the MLP algorithm outperformed other models. It achieved the lowest MAE, MSE, and RMSE values, indicating high precision in droplet volume predictions. Additionally, the MLP’s R-squared value was close to 1, reflecting a strong correlation between the predicted and actual droplet volumes. This highlights the MLP’s robustness in handling the complexities of droplet bioprinting parameter predictions.

Regarding computation time, the DT algorithm exhibited the shortest training and testing times due to its simpler architecture relatively compared to DL ^43^. This makes it an attractive option for scenarios where rapid computation is essential. Nonetheless, the MLP also demonstrated efficient computational performance, with training times under 4 seconds and testing times under 2 seconds, suggesting that it can be effectively used in real-time bioprinting applications without significant delays.

Overall, this study highlights the potential of ML and DL algorithms in optimizing bioprinting processes. The MLP, with its superior prediction accuracy and reasonable computation time, offers a balanced approach to enhancing bioprinting precision. Meanwhile, the DT algorithm provides an alternative when computation speed is the priority. These insights contribute to the advancement of bioprinting technologies, ultimately aiding in creating high-quality, reproducible bioprinted constructs for applications in tissue engineering and regenerative medicine.

## Methods

### Synthesis and preparation of bioinks

GelMA-based bioink has been demonstrated to enhance cell viability and improve printing quality during the extrusion-based bioprinting process ^44^. The synthesis of GelMA was conducted according to a previously reported protocol ^44^. In summary, the outlined procedure involves the dissolution of 5g of porcine skin-derived powdered gelatin (Type A, Bloom strength 300, Sigma-Aldrich, St. Louis, MO, USA) in 50 mL of distilled water. Upon achieving complete dissolution at a temperature of 50°C, 10 mL of glycidyl methacrylate (Sigma-Aldrich, St. Louis, MO, USA) is gradually added dropwise into the dissolved gelatin solution. The resultant solution is maintained for 12 hours at 50°C with continuous stirring at 750 RPM. Subsequently, the solution is transferred into a dialysis tube with a molecular weight cutoff of 12-14 kDa (Fisher Scientific, Waltham, MA, USA). Over the subsequent three days, the water in the dialysis tube tank is replaced twice daily. Following this period, the resulting solution is frozen at −80°C for 24 hours, followed by lyophilization at −84°C for three days to obtain dried GelMA foam. The synthesized GelMA is then stored at −20°C for future use. To prepare the GelMA-Alginate hydrogel, a stock solution of 5% (w/v) GelMA in PBS (VWR, Radnor, PA, USA) containing 0.2% lithium phenyl-2,4,6-trimethylbenzoylphosphinate (LAP) photoinitiator (Sigma-Aldrich, St. Louis, MO, USA) (referred to as stock 1). The presence of alginate can support the stability of hydrogel precursor when bioprinting ^45^. The increased viscosity of the hydrogel precurosr due to the addition of alginate is helpful especially when bioprinting very small droplets, ranging from 0.1 to 2 µL. Alginate solutions with concentrations of 0.5% and 1% (w/v) in stock 1 were prepared, and vigorous agitation was employed for 3 hours for complete dissolution.

Green fluorescent protein (GFP) tagged NIH 3T3 fibroblast cells (ATCC, Manassas, VA, USA) were utilized for bioprinting optimization and in vitro experiments. Cells were maintained in Dulbecco’s Modified Eagle Medium (VWR, Radnor, PA, USA) supplemented with 10% Fetal Bovine Serum (VWR, Radnor, PA, USA) and 1% Penicillin-Streptomycin (VWR, Radnor, PA, USA). Culture conditions were maintained at 37°C and 5% CO_2_, with growth medium replacement every 48 hours. Upon reaching 90% confluency, cells were harvested by incubation in trypsin-EDTA solution (VWR, Radnor, PA, USA) for 5 minutes followed by centrifuging for 3 minutes at 1500 RPM. For optimization and in vitro characterization, the harvested cells were suspended in two bioink formulations: 5% GelMA with 1% Alginate (hereinafter referred to as 5G1A) and 5% GelMA with 0.5% Alginate (5G0.5A). The hydrogel characterization proceeded with 5% GelMA (5G) and 5% GelMA with 2% Algiante (5G2A) as well.

### Bioink characterization

Rheological property characterization was conducted using a rheometer (MCR 302e, Anton Paar, Graz, Austria) equipped with a Peltier plate for the 5G1A and 5G0.5A materials used in the machine learning training and 5G and 5G2A as the control groups. A stainless-steel parallel plate with a diameter of 25 mm and a gap distance of 0.5 mm at room temperature was utilized for all experiments. Rheological tests assessed the flow characteristics and viscoelastic behavior pertinent to the bioprinting process.

To verify the photocrosslinking kinetics, a transparent glass plate was integrated into the experimental setup, allowing for the illumination of the hydrogel with 405 nm light source from beneath the 25 mm parallel plate geometry. The initial 60-second period with the light off establishes the baseline viscoelastic properties of the uncrosslinked materials. Upon light activation at 60 seconds, the evolution of storage modulus (G’) and loss modulus (G’’) provides insight into the hydrogels’ crosslinking kinetics and final mechanical properties. Oscillatory measurements were conducted using a 1% shear strain and a frequency of 1 Hz. Concurrently, both the storage and loss moduli for the photocrosslinking bioink were recorded.

The transparency of samples printed in the shape of a disc was investigated to enhance the efficiency of the light permeability during the photocrosslinking process. In addition, the transparent sample can facilitate the optical observation of cells and their surrounding environment within the hydrogel. To assess the transparency, crosslinked samples were prepared in the form of disks, each with a diameter of 8 mm and a height of 6 mm, and placed on a sheet of paper bearing the university logo.

To determine the viscosity of materials 5G1A and 5G0.5A, the zero-shear viscosity was measured by setting the shear rate to 1 s^−1^. The mechanical properties of photocrosslinked hydrogel were evaluated by measuring the compressive elastic modulus, which reflects the stiffness of its microstructure. Hydrogel precursors were dispensed into 8.5 mm diameter and 3.5 mm height molds, with each disk receiving 300 μL. After crosslinking with 405 nm light for 1 minute, the samples were immediately tested for compressive modulus. A vertical-axis micromechanical testing machine (ESM 301L, Mark-10, USA) equipped with a custom-made flat-ended rigid cylinder (12.5 mm in diameter) was utilized to conduct compression tests. This equipment recorded force-displacement measurements during indentation. Force-displacement plots were generated by subjecting the hydrogel surface to a 50% displacement at a 10 mm/min speed. Utilizing the initial dimensions of the hydrogel samples and custom MATLAB code, the compression modulus was calculated based on the slope within the initial 10% strain region, known as the linear region.

Evaluating water absorption capacity in crosslinked hydrogel samples requires assessing their swelling ratio. Initially, 300 μL of pre-polymer solutions were loaded into molds with an 8 mm diameter. These solutions were then crosslinked under 405 nm light for 1 minute. After crosslinking, the hydrogels were immersed in phosphate-buffered saline (PBS) and kept in a 37°C environment for 24 hours to ensure complete hydration. Upon achieving equilibrium swelling, the weight of each sample in its hydrated state (W_w_) was accurately recorded. Subsequently, the hydrated hydrogel samples were frozen in a −80°C freezer and subjected to lyophilization for three days to determine their dry weight (W_d_). Finally, **Equation (2)** was utilized to calculate the swelling ratio, with the entire process being repeated three times for consistency.

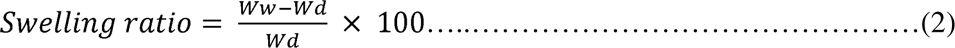

### Development of a 3D bioprinting platform specialized in machine learning-driven droplet optimization

A fully customizable 3D bioprinting system was developed by incorporating additional components to facilitate the high-throughput dataset generation essential for training ML algorithms. The bioprinting process is controlled by pre-defined G-code instructions (provided in supplementary) that direct the bioink dispensing and print bed movement based on droplet volume and spacing between each droplet’s requirements, ensuring precise deposition and preventing droplet merging. Another advantageous feature of this system is its micro-dropletization capability, i.e., producing droplets with an average diameter of approximately 100 μm. This functionality is particularly valuable for constructing single-cell arrays, allowing for the precise delivery of individual cells to specific locations. The system’s efficiency is demonstrated by its ability to rapidly and accurately bioprint 60 droplets, each with a 4 μL volume, on a coated hydrophobic glass slide (VWR, Radnor, PA, USA) (**Supplementary Movie S7**).

Accurate measurement of droplet volume necessitated the determination of both diameter and contact angles. The diameter was determined using a calibrated pixel-to-length ratio method. This approach established the length represented by each pixel, providing a reliable basis for precise diameter calculation and ensuring accuracy in the quantitative analysis of droplet dimensions.

### Droplet volume calculation and image processing

The calculation of contact angles was performed through image processing. In the methodology proposed in this work, the raw images undergo a comprehensive series of three distinct image processing stages. Initially, the recorded multi-channel images are converted into grayscale, thereby optimizing memory utilization by integrating the three red, green, and blue channels into a single-channel grayscale matrix. With a specific focus on contrast and brightness parameters, this grayscale conversion facilitates the extraction of clearer droplet features, enhancing visual discernibility. The second stage involves Canny edge detection processing. Employing a Gaussian filter mitigates noise within the image, leading to a more precise delineation of droplet edges. Subsequently, the images undergo conversion to a binary format. Binary images, characterized by pixels with only two values, streamline subsequent image-processing tasks. The relevant area is represented in black to accentuate the droplet pattern within the image, while the remaining portions are depicted in white. This deliberate binary representation effectively highlights the droplet within the image. In the final stage, all processed images, encompassing grayscale, Canny edge detection, and binary representations, are combined and saved as the final combined image.

### Bioprinting preparations and parameters

The silane-coated glass substrate traveled along the x-axis to accommodate multiple cell-laden droplets at uniform spacing. These deposited droplets were then placed under the 405 nm lamp to polymerize the photo-crosslinkable biomaterials. Printing parameters and corresponding droplet volume were used as input datasets when training with the ML and DL algorithms (Table 4). Two bioink formulations with different viscosities were utilized: 5G0.5A (10.03 mPa·s) and 5G1A (24.66 mPa·s). The bioprinting process employed three conical needles with varying inner diameters of dispensing nozzles. Printing parameters were systematically varied, with printing times set at 0.05, 0.1, and 0.15 seconds and printing pressures adjusted to 1.5 and 2 psi, respectively. The cell-laden biomaterials were prepared in 3 mL volumes, with two different cell concentrations: 2.8×10^6^ and 5.6×10^6^ cells/3 mL.

### Machine learning (ML)

In this study, we implemented the supervised learning domain, a fundamental paradigm in ML that utilizes labeled datasets. Within this framework, our research focuses on regression models because the primary objective of this study was to develop models capable of predicting continuous output values based on corresponding input data.

### Decision tree (DT)

The decision tree (DT) concept encompasses a structure reminiscent of a combination of trees and branches, with its principal characteristic being its swift operation even on intricate datasets ^46^. The DT partitions data based on specific criteria or questions, and the algorithm’s efficacy relies on generating optimized questions. Initiated by the classification and regression tree algorithm, which commences from the root node representing the initial depth and proceeds to form a binary tree, we divided the data into two regions for each node branch ^47,48^. Each branch comprises a rule node at the midpoint and a leaf node at its terminus, providing the sought-after answer. While a greater number of rule nodes enhance learning for prediction, the algorithm concurrently introduces complexity, potentially leading to overfitting. We trained a DT algorithm using 10-fold cross-validation with max_depth values ranging from 1 to 19. For each depth, the model was trained using 10 different training and test datasets. We recorded the values of train_mae (training dataset mean abolute error), val_mae (validation dataset mean absolute error), train_rmse (training dataset root mean square error), val_rmse (validation dataset root mean square error), train_r2 (training dataset R-squared), val_r2 (validation dataset R-sqaured), train_mse (training dataset mean squared error), and val_mse (validation dataset mean squared error). Analysis of these results indicated that a max_depth of 7 provides the optimal balance, effectively avoiding overfitting.

DT finds application in both classification and regression scenarios. Three metrics—Gini index, entropy, and classification error—are employed to measure data impurity in a binary DT ^47^. A lower Gini index signifies greater data uniformity, and the method yielding the highest information gain value is selected to derive results. Impurity serves as a classification standard, facilitating the creation of branches with lower impurity by distinctly distinguishing between classes collecting identical objects. Conversely, in the DT tailored for regression models, impurity is gauged using the mean square error (MSE) ^49^.

### Random forest (RF)

In RF, five primary types of hyperparameters (e.g., n_estimators, criterion, max_depth, min_samples_split, and min_samples_leaf) exist (**Supplementary Table S3**). The process of hyperparameter tuning is essential for implementing an optimal training model ^50^. In the process of analyzing data using a RF model, we aimed to determine the optimal number of estimators (n_estimators) by experimenting with various values ranging from 1 to 19. To achieve this, we employed 10-fold cross-validation, which involved splitting the data into 10 distinct training and test datasets for each number of estimators. This approach allowed us to comprehensively evaluate the model’s performance and identify the most appropriate number of trees in the forest while avoiding overfitting.

By comparing the performance metrics across different numbers of estimators, we identified that using 10 estimators provided the best balance between training and validation performance. At this number, the model exhibited robust performance on both datasets, signifying that it effectively captured the underlying patterns in the data without overfitting. Based on this analysis, we concluded that the optimal number of estimators for our RF model is 10. This number allows the model to maximize its predictive performance on new data while maintaining a balance between complexity and generalization.

While optimal conditions for these five hyperparameters can be conducted manually, this study employed an automated training method with optimized hyperparameter combinations. RandomizedSearchCV randomly selects a specified number of combinations from a predefined hyperparameter search space and outputs the combination yielding the best score ^51^. Conversely, GridSearchCV exhaustively explores all conceivable combinations of hyperparameter values. In this paper, GridSearchCV was used as it showed better performance ^52^.

### Polynomial regression (PR)

In contrast to linear regression (**Equation 3**), which establishes a linear association between a single independent variable and a dependent variable, and multiple linear regression (**Equation 4**), which involves multiple independent variables and a single dependent variable, polynomial regression (PR) analysis utilizes a polynomial function ^53^. This approach captures non-linear relationships between independent and dependent variables, leveraging polynomial functions to discern intricate patterns within the data and facilitate predictive modeling. In PR, the hyperparameter known as the degree of polynomial is used to ascertain the highest dimension utilized in the polynomial (**Equation 5**) ^54,55^.

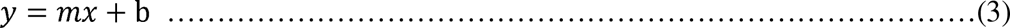

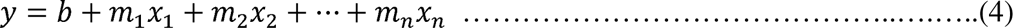

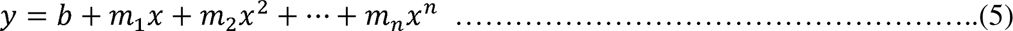

In analyzing data with a PR model, we tested polynomial degrees from 1 to 9. We found that performance improved with increasing degrees on the training dataset but worsened on the validation dataset beyond a certain point, indicating overfitting. By comparing performance metrics, we determined that a polynomial degree of 7 offered the best balance between training and validation performance, effectively capturing data patterns without overfitting. Thus, we concluded that a polynomial degree of 7 is optimal for maximizing predictive performance while maintaining a balance between complexity and generalization.

### Deep learning (DL)

DL is a subset of ML, defined by utilizing multi-layer neural networks that consist of several layers, including an input layer, hidden layers, and an output layer. Often referred to as artificial neural networks, these multi-layer neural networks form the basis of DL^56,57^.

### Multi-layer perception (MLP)

To determine the optimal number of epochs for our MLP model, we tested epoch counts from 10 to 100 in increments of 10 using 10-fold cross-validation. This approach involved splitting the data into 10 training and test sets for each epoch count, allowing for a thorough evaluation while preventing overfitting. We found that performance improved with more epochs on the training dataset, but beyond a certain point, validation performance stabilized, indicating effective learning without significant overfitting. Comparing metrics across epoch counts, 100 epochs provided the best balance between training and validation performance. Thus, 100 epochs is optimal for maximizing predictive performance while balancing complexity and generalization.

### Long short-term memory (LSTM)

LSTM, as a type of recurrent neural network, is commonly employed in DL for processing sequential or time-dependent data ^58^. Unlike feedforward neural networks such as MLP, which are well-suited for non-sequential data, LSTM excels at capturing patterns in sequences. In the process of analyzing data using an LSTM model, we aimed to determine the optimal number of epochs by experimenting with various epochs ranging from 10 to 100 in increments of 10. We used 10-fold cross-validation to evaluate the model’s performance across different epoch counts. This method involved splitting the data into 10 training and test sets per epoch count, allowing us to assess model performance comprehensively and prevent overfitting. We found that while performance on the training dataset improved with more epochs, it began to decline on the validation dataset after a certain point, indicating overfitting. Comparing performance metrics, we determined that 100 epochs struck the best balance between training and validation performance, effectively capturing data patterns without overfitting. Therefore, we concluded that 100 epochs is the optimal number for our LSTM model, maximizing predictive performance while balancing complexity and generalization.

### Analysis of the dataset

Each model underwent a dataset partitioning process, wherein the entire dataset, comprising a total of 1758 instances, was randomly partitioned into two sets: a training set, constituting 70% of the data, and a testing set, comprising the remaining 30% ^60^. Even within the 70% training set, 10-fold cross-validation was performed, wherein the dataset was partitioned into ten mutually exclusive subsets. Cross-validation was utilized to assess the performance of the models systematically and enhance their generalization capabilities ^61^. The model training and evaluation processes were iteratively executed ten times, each time utilizing a different subset for testing while the remaining data was used for training. This iterative procedure resulted in ten individual performance scores, which were then aggregated through averaging to yield the final comprehensive evaluation of model performance. The ultimate performance metric was derived from the outcomes of the testing set. Image preprocessing, data analysis, and training of the machine and DL were implemented using Python version 3.12.1 with TensorFlow and sci-kit-learn (Provided at GitHub). Detailed information about the model used and tuning parameters is provided in the following sections.

### Optimization of the hyperparameters

Hyperparameters serve as configurations to effectively control and adjust the learning process of a model before its training ^62^. The process of tuning hyperparameters holds significant importance, impacting the structure of the model, the learning procedure, and the optimization method employed. In the context of this study, distinct hyperparameters were employed for each of the algorithms. The Max Depth hyperparameter was fine-tuned for the DT algorithm to determine the maximum tree depth ^63^. RF underwent tuning using five specified hyperparameters (**Supplementary Table S3**). In the case of PR, the polynomial degree hyperparameter was manipulated to establish the polynomial’s degree. Conversely, in DL models such as MLP and LSTM, the optimal hyperparameters are autonomously selected from the data within the model itself, eliminating the need for prior hyperparameter tuning efforts.

### Evaluation metrics for comparative analysis between machine learning algorithms

The regression-based ML and DL models presented in this study were compared and analyzed using four evaluation criteria. First, the mean absolute error (MAE) measures the absolute difference between predicted and actual values ^35^. A lower MAE suggests that the model’s predictions are considered closer to the actual values. Mean Squared Error (MSE) measures the squared difference between predicted and actual values. MSE involves squaring each prediction error, summing up these squared values, and then dividing by the total number of data points to calculate the mean. Consequently, it always produces non-negative values and assigns higher weights to larger errors. A lower MSE indicates that the model’s predictions are considered closer to the actual values. Root Mean Squared Error (RMSE) is an advanced form of MSE obtained by taking the square root of the MSE values. It allows us to calculate the average magnitude of prediction errors, and a smaller RMSE value is also interpreted as the model’s predictions being closer to the actual values. R-squared is a metric used to assess the goodness of fit in a regression model, indicating how well the model aligns with the provided data on a scale between 0 and 1 ^35^. A value closer to 1 signifies a well-trained model, demonstrating high suitability to the given data. The higher the R-squared value, the better the model is considered to explain the data. However, it is crucial to note that R-squared can also approach 1 in cases of overfitting. Therefore, using other evaluation metrics in conjunction with R-squared is advisable.

### Statistical analysis

To evaluate the performance differences among three ML and two DL algorithms and other relevant factors, both one-way and two-way analyses of variance (ANOVA) were conducted in Minitab 20 (Minitab, LLC., PA, USA). The null hypothesis posited that all algorithms exhibited equivalent performance. Rejection of this null hypothesis would indicate statistically significant differences in prediction performance among the groups. The ANOVA was followed by Tukey’s Honestly Significant Difference (HSD) test as a post-hoc analysis. Tukey’s HSD allows for identifying specific algorithms that differ significantly in their mean performance. The evaluation framework employed a 10-fold cross-validation approach, necessitating ten iterations of the analysis. This robust methodology ensures the reliability and generalizability of the performance comparisons across the ML algorithms under investigation.

## Conclusions

We introduce a highly efficient bioprinting platform capable of generating high-throughput cellular droplets, integrated with an artificial intelligence (AI)-based optimization system. Our innovative approach leverages the power of AI to significantly enhance the bioprinting process, which traditionally depends on time-consuming trial-and-error methods to achieve optimal droplet volumes for organoids and cellular droplet bioprinting. Droplet volumes were calculated by analyzing droplet diameters and contact angles from high-resolution images, ensuring unparalleled accuracy in our predictions. By utilizing three modified and optimized ML algorithms and two DL algorithms, we have developed predictive models that accurately determine cell-laden droplet volumes based on a comprehensive set of bioprinting parameters. These models have shown exceptional proficiency in learning the complex relationships between input parameters and droplet volume outcomes. The top-performing model demonstrated remarkable prediction accuracy and impressively short training and testing times, marking a significant advancement in the efficiency and effectiveness of bioprinting applications. We employed four regression model evaluation metrics to assess the results. Each metric provides valuable insights depending on the study’s objectives and the nature of the data. The RMSE evaluation index shows MLP and LSTM as the most accurate regression model. In the R-squared evaluation index that shows the volatility of data, DT, RF, PR, and MLP all show the best results with values above 0.95. In terms of computational time, the ML algorithm showed a much faster speed than the DL algorithm. This innovative approach streamlines the optimization process and significantly reduces bioink wastage and time, addressing a major challenge in the field. In addition, our innovative approach represents a significant leap forward in the field of bioprinting. Combining cutting-edge ML techniques with advanced bioprinting technology opens new tissue engineering and regenerative medicine frontiers. This breakthrough promises to accelerate research, improve therapeutic outcomes, and ultimately contribute to better health and well-being for countless individuals worldwide.

## Acknowledgments

This work was supported by a Natural Sciences and Engineering Research Council of Canada (NSERC) Discovery Grant (RGPIN-2020-04559) and the Canada Foundation for Innovation John R. Evans Leaders Opportunity Fund. J.S. was supported by the NSERC Vanier Canada Graduate Scholarship and Alberta Innovate Graduate Student Scholarship.

## Contributions

The project and concept were conceived by J.S. and K.K. J.S. conducted most of the experiments. M.K. contributed to data processing and developed the web user interface. K.H. and H.K. assisted in modifying the 3D bioprinter system. J.S. wrote the final manuscript with contributions from all authors. All authors discussed the results and provided comments on the manuscript.

## Ethics declarations

The authors declare no competing interests.

## Data availability

All relevant data supporting this study’s key findings, as well as the raw image data generated in this study, are available within the paper and its Supplementary Information or from the corresponding author upon reasonable request. All G-codes used for bioprinting are included in the Supplementary Information. Code for image processing, machine learning, and deep learning is publicly available on GitHub at https://github.com/ryanKang-programmer/dropletMeasurement.

