## Supplementary Materials for "Machine Learning Driven Optimization for High Precision Cellular Droplet Bioprinting"

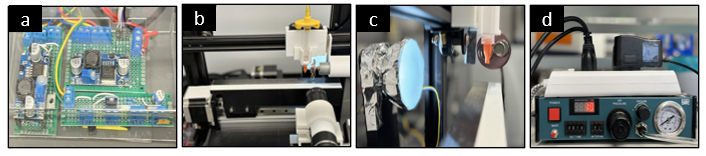

**Supplementary Fig. S1:** Components of the high-throughput printing and imaging system. **a** Custom-designed circuit board for system control and data acquisition. **b** Motorized stage assembly for precise positioning and high-throughput image capture. **c** Optimized LED lighting configuration for consistent illumination across samples. **d** Pneumatic dispenser unit for accurate and controlled sample deposition.

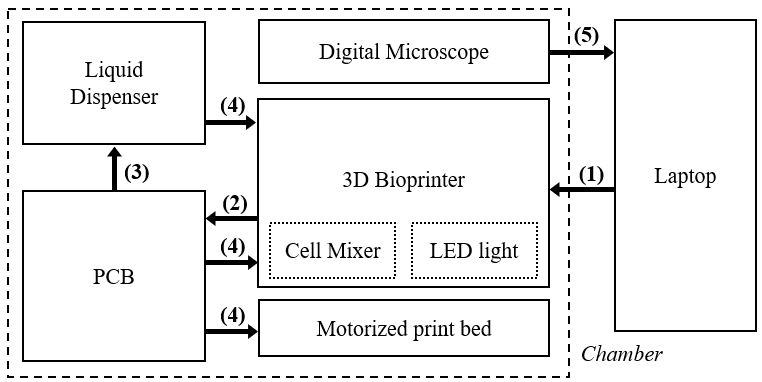

**Supplementary Fig. S2:** A comprehensive signal diagram illustrating the workflow of a whole bioprinting platform. The setup includes a laptop connected to a 3D bioprinter and a modified printed circuit board (PCB) integrated with a liquid dispenser. This interconnected system facilitates precise droplet dispensing and subsequent image collection, highlighting the seamless coordination between hardware components and software control for efficient bioprinting operations.

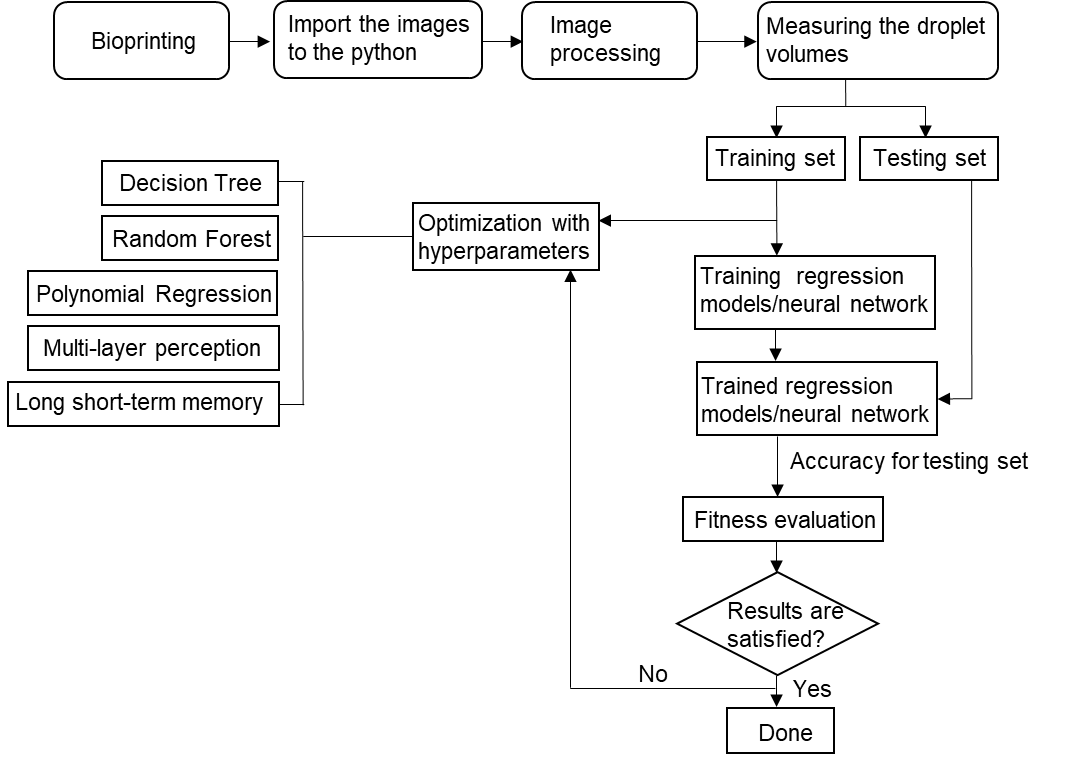

**Supplementary Fig. S3:** Integrated framework for bioprinting parameter optimization using machine learning and deep learning algorithms.

**Supplementary Table S1.** Comparison between machine learning and deep learning.

|  | **Machine learning** | **Deep learning** |
| --- | --- | --- |
| **Data size** | Efficient with modest-sized datasets and suited for tasks of moderate complexity. | Excels on large and intricate datasets, suitable for high-dimensional and complex data. |
| **Complexity of model** | General and shallow model in a concise and formal tone. | Utilization of a multilayer neural network structure and complex model. |
| **Hardware and optimization** | Not specifically dependent on hardware, applicable to various architectures | Specific hardware (GPU, TPU) requirements and optimization techniques are necessary for efficient training. |
| **Batch processing** | Processing data in small batches | It is generally more suitable for processing in bulk batches to efficiently handle large-scale datasets. |
| **Hyperparameter adjustment** | Hyperparameters are typically manually tuned based on domain knowledge and experimentation. | Adjusting more hyperparameters requires utilizing automated methods for hyperparameter tuning. |

**Supplementary Table S2.** Statistical analysis of bioprinting parameter weights on droplet volume using the Tukey method with a 95% confidence interval

| **Parameter** | **N** | **Mean** | **Grouping** |
| --- | --- | --- | --- |
| Dispensing time | 5 | 0.387772 | A |
| Nozzle gauge | 5 | 0.257286 | B |
| Printing pressure | 5 | 0.196495 | C |
| Cell concentration | 5 | 0.114062 | D |
| Viscosity | 5 | 0.044384 | E |

*Means that do not share a letter are significantly different.*

**Supplementary Table S3.** Types of hyperparameters utilized in optimization before algorithm training: their functions and roles

| **Hyperparameter** | **Function** |
| --- | --- |
| n_estimators | number of trees |
| criterion | impurity indicators (gini, entropy, log_loss) |
| max_depth | maximum depth of tree |
| mim_samples_split | minimum number of samples required to split an internal node |
| min_samples_leaf | minimum number of samples a leaf node must have |

**Supplementary Movie S1**. High-throughput cellular microarray 3d bioprinting

**Supplementary Movie S2**. Bioprinting workflow on the integrated platform

**Supplementary Movie S3**. Bioprinting of droplets with a minimum volume of 0.1 µL

**Supplementary Movie S4**. Bioprinting of 60 droplets on a single glass slide

**Supplementary Movie S5.** Crosslinking with 405 nm Blue Light Post-Printing

**Supplementary Movie S6.** Cell Stirring system in syringe to prevent cell sedimentation

**Supplementary Movie S7**. Day 3 of the time-lapse movie of GFP-tagged 3T3 fibroblast cells encapsulated in 5% Gelma and 2% Alginate bioink droplets (~500 µm thickness) across different focal planes

**G-code for bioprinting**

G92 X0 Y0 E0

M302 S0 ; cold extrusion at any temperature

M104 S60 ; set temperature to 60degC

G91

;start

;M104

G01 X0 Y0 Z-5 F3000

M106

G4 P2000

M107

G01 X0 Y0 Z5 F3000

G1 E-30 F800 ; move extruder motor/platform motor.

G92 E0

M104 S60

G4 P2000

G01 X0 Y0 Z-5 F3000

M106

G4 P2000

M107

G01 X0 Y0 Z5 F3000

G1 E-30 F800

G92 E0

M104 S60

G4 P2000

G01 X0 Y0 Z-5 F3000

M106

G4 P2000

M107

G01 X0 Y0 Z5 F3000

G1 E-30 F800

G92 E0

M104 S60

G01 X0 Y0 Z-5 F3000

M106

G4 P2000

M107

G01 X0 Y0 Z5 F3000

G1 E-30 F800

G92 E0

M104 S60

G01 X0 Y0 Z-5 F3000

M106

G4 P2000

M107

G01 X0 Y0 Z5 F3000

G1 E-30 F800

G92 E0

M104 S60

G01 X0 Y0 Z-5 F3000

M106

G4 P2000

M107

G01 X0 Y0 Z5 F3000

G1 E-30 F800

G92 E0

M104 S60

G01 X0 Y0 Z-5 F3000

M106

G4 P2000

M107

G01 X0 Y0 Z5 F3000

G1 E-30 F800

G92 E0

M104 S60

G01 X0 Y0 Z-5 F3000

M106

G4 P2000

M107

G01 X0 Y0 Z5 F3000

G1 E-30 F800

G92 E0

M104 S60

G01 X0 Y0 Z-5 F3000

M106

G4 P2000

M107

G01 X0 Y0 Z5 F3000

G1 E-30 F800

G92 E0

M104 S60

G01 X0 Y0 Z-5 F3000

M106

G4 P2000

M107

G01 X0 Y0 Z5 F3000

G1 E-30 F800

G92 E0

M104 S60

G01 X0 Y0 Z-5 F3000

M106

G4 P2000

M107

G01 X0 Y0 Z5 F3000

G1 E-30 F800

G92 E0

M104 S60

G01 X0 Y0 Z-5 F3000

M106

G4 P2000

M107

G01 X0 Y0 Z5 F3000

G1 E-30 F800

G92 E0

M104 S60

G01 X0 Y0 Z-5 F3000

M106

G4 P2000

M107

G01 X0 Y0 Z5 F3000

G1 E-30 F800

G92 E0

M104 S60

G01 X0 Y0 Z-5 F3000

M106

G4 P2000

M107

G01 X0 Y0 Z5 F3000

G1 E-30 F800

G92 E0

M104 S60

G01 X0 Y0 Z-5 F3000

M106

G4 P2000

M107

G01 X0 Y0 Z5 F3000

G1 E-30 F800

G92 E0

M104 S60

G01 X0 Y0 Z-5 F3000

M106

G4 P2000

M107

G01 X0 Y0 Z5 F3000

G1 E-30 F800

G92 E0

M104 S60

G01 X0 Y0 Z-5 F3000

M106

G4 P2000

M107

G01 X0 Y0 Z5 F3000

G1 E-30 F800

G92 E0

M104 S60

G01 X0 Y0 Z-5 F3000

M106

G4 P2000

M107

G01 X0 Y0 Z5 F3000

G1 E-30 F800

G92 E0

M104 S60

G01 X0 Y0 Z-5 F3000

M106

G4 P2000

M107

G01 X0 Y0 Z5 F3000

G1 E-30 F800

G92 E0

M104 S60

G01 X0 Y0 Z-5 F3000

M106

G4 P2000

M107

G01 X0 Y0 Z5 F3000

G1 E-30 F800

G92 E0

M104 S60

G01 X0 Y0 Z-5 F3000

M106

G4 P2000

M107

G01 X0 Y0 Z5 F3000

G1 E-30 F800

G92 E0

M104 S60

G01 X0 Y0 Z-5 F3000

M106

G4 P2000

M107

G01 X0 Y0 Z5 F3000

G1 E-30 F800

G92 E0

M104 S60

G01 X0 Y0 Z-5 F3000

M106

G4 P2000

M107

G01 X0 Y0 Z5 F3000

G1 E-30 F800

G92 E0

M104 S60

G01 X0 Y0 Z-5 F3000

M106

G4 P2000

M107

G01 X0 Y0 Z5 F3000

G1 E-30 F800

G92 E0

M104 S60

G01 X0 Y0 Z-5 F3000

M106

G4 P2000

M107

G01 X0 Y0 Z5 F3000

G1 E-30 F800

G92 E0

M104 S60

G01 X0 Y0 Z-5 F3000

M106

G4 P2000

M107

G01 X0 Y0 Z5 F3000

G1 E-30 F800

G92 E0

M104 S60

G01 X0 Y0 Z-5 F3000

M106

G4 P2000

M107

G01 X0 Y0 Z5 F3000

G1 E-30 F800

G92 E0

M104 S60

G01 X0 Y0 Z-5 F3000

M106

G4 P2000

M107

G01 X0 Y0 Z5 F3000

G1 E-30 F800

G92 E0

M104 S60

G01 X0 Y0 Z-5 F3000

M106

G4 P2000

M107

G01 X0 Y0 Z5 F3000

G1 E-30 F800

G92 E0

M104 S60

G1 E450 F800 ; move extruder motor/platform motor.

M104 S60
